# *Streptococcus agalactiae npx* is required for survival in human placental macrophages and full virulence in a model of ascending vaginal infection during pregnancy

**DOI:** 10.1101/2022.10.20.513045

**Authors:** Jacky Lu, Rebecca E. Moore, Sabrina K. Spicer, Ryan S. Doster, Miriam A. Guevara, Jamisha D. Francis, Kristen N. Noble, Lisa M. Rogers, Julie A. Talbert, Michelle L. Korir, Steven D. Townsend, David M. Aronoff, Shannon D. Manning, Jennifer A. Gaddy

## Abstract

*Streptococcus agalactiae*, also known as Group B *Streptococcus* (GBS), is a Gram-positive encapsulated bacterium that colonizes the gastrointestinal tract of 30-50% of humans. GBS causes invasive infection during pregnancy that can lead to chorioamnionitis, funisitis, preterm prelabor rupture of membranes (PPROM), preterm birth, neonatal sepsis, and maternal and fetal demise. Upon infecting the host, GBS encounters sentinel innate immune cells, such as macrophages, within reproductive tissues. Once phagocytosed by macrophages, GBS upregulates expression of the gene, *npx*, which encodes a NADH peroxidase. GBS mutants with a *npx* deletion (Δ*npx*) are exquisitely sensitive to reactive oxygen stress. Furthermore, we have shown that *npx* is required for GBS survival in both THP-1 and placental macrophages. In an *in vivo* murine model of ascending GBS vaginal infection during pregnancy, *npx* is required for invasion of reproductive tissues and is critical for inducing disease progression including PPROM and preterm birth. Reproductive tissue cytokine production was also significantly diminished in Δ*npx* infected animals compared to those infected with wild type (WT)-GBS. Complementation *in trans* reversed this phenotype, indicating *npx* is critical for GBS survival and initiation of proinflammatory signaling in the gravid host.

## Introduction

Preterm deliveries occur at less than 37 weeks gestation (1). Despite advancing knowledge of risk factors and introduction of public health and medical interventions to reduce the occurrence, the rate of preterm birth in the United States and other developed countries still hovers between 5 and 9% (2, 3). Preterm birth accounts for 75% of perinatal mortality and over 50% of the long-term morbidity (4). While most preterm babies survive, these infants are at increased risk of neurodevelopmental impairments, respiratory conditions, and gastrointestinal complications (5). Intrauterine infection may account for at least 25-40% of preterm births (6).

*Streptococcus agalactiae*, or Group B *Streptococcus* (GBS), is an encapsulated Gram-positive bacterium that colonizes the urogenital tract and lower gastrointestinal tract in 30% of healthy adults (7). Although GBS is a common member of the intestinal microbiota, it can cause invasive infections during pregnancy, leading to sepsis or meningitis in the neonate (8). Indeed, GBS is a leading cause of adverse pregnancy and neonatal outcomes such as stillbirth, chorioamnionitis, preterm birth, and neonatal sepsis (9); up to 25% of invasive GBS infections during pregnancy end in stillbirth or spontaneous abortion (10). To prevent neonatal infection, the Centers for Disease Control and Prevention (CDC) recommends screening mothers for GBS late in the third trimester and administering antibiotic therapy to those who test positive during labor (11). There is concern, however, that antibiotic exposure could alter the infant’s developing microbiome, which may contribute to life-long consequences (8), and intrapartum antibiotic prophylaxis does not prevent late-onset disease, stillbirth, or preterm births (12). Consequently, GBS remains the leading infectious cause of morbidity and mortality among neonates in the United States (13).

GBS pathogenesis begins with adherence to vaginal epithelial cells (8). For successful colonization, the bacteria can form biofilm structures to evade the immune system (14). Following vaginal colonization, GBS can ascend above the cervical os, through as-yet undefined mechanisms, and traverse the fetal membranes, causing fetal infection (15). The inflammation of extraplacental (“fetal”) membranes in response to GBS infection is termed chorioamnionitis (16).

Because the human immune system has evolved to protect against pathogens, this paradigm is more complex during pregnancy as the system must defend the gravid uterus against infection and maintain immunologic tolerance to the semi-allogenic fetus. This careful balance must be maintained to prevent harm to the mother and fetus. Consequently, the immune response during pregnancy is characterized by dynamic modifications in the maternal and fetal tissues reliant on the stage of pregnancy (17, 18). Bacteria commonly cause intrauterine infection, triggering a proinflammatory response originating in the decidua by activation of pattern recognition receptors, which may result in preterm birth (19, 20). Previous mouse studies have demonstrated that innate immune signaling is sufficient to instigate adverse pregnancy outcomes (18). The presence of proinflammatory cytokines, IL-1β, IL-6, IL-8, and TNF-α, in the amnionic fluid or in cervico-vaginal lavage in patients are indicative of the onset of preterm labor (21–23). A variety of leukocytes responsible for cytokine production are present in the reproductive tissues, including maternal natural killer cells, dendritic cells, macrophages, and lymphocytes (24). In particular, macrophages represent a predominate subset of human leukocytes that serve as antigen presenting cells (APC) in the decidua, comprising 20-25% of all decidual leukocytes (25).

Placental macrophages (PMs) represent a mixed population of maternal and fetal-derived cells (26) that are thought to play critical roles in placental invasion, angiogenesis, tissue modeling, and development (27, 28). Recently, studies have demonstrated that PMs defend against invading bacterial pathogens by release of macrophage extracellular traps (29).

Upon phagocytosis by a macrophage, bacterial pathogens are trapped in a phagosome, which is a highly oxidative environment (33). Additionally, cells generate reactive oxygen species (ROS) as metabolic byproducts. The amount of ROS produced by the NOX2 NADPH oxidase in macrophages is significantly higher under conditions of infection compared to resting states; hence, stimulating oxidative stress to kill invading pathogens is a critical pathway for innate immunity (33). ROS can damage macromolecules including lipids, proteins, and nucleic acids, ultimately leading to cell death (34). Bacterial pathogens have evolved strategies to survive in highly oxidative environments such as producing antioxidants or enzymes that can inactivate and detoxify ROS (35). GBS produces several products to help circumnavigate ROS stress during infection including superoxide dismutase, which converts superoxide to H_2_O_2_ and O_2_ (36). GBS also produces glutathione and a carotenoid pigment which protect against oxidative damage (37, 38). A previous study revealed that GBS *npx* encodes an NADH peroxidase, which is critical for detoxification and resistance to peroxide stress and survival within THP-1 macrophage-like cells (39). Deletion of the *npx* locus in the Δ*npx* mutant resulted in attenuated ability to detoxify peroxide; a result that was reversed via genetic complementation *in trans* (39).

During ascending pregnancy infections, one of the first immune cells GBS encounters at the maternal-fetal interface are placental macrophages. We hypothesized that *npx* is also required for GBS survival within PMs and to cause disease progression during pregnancy. To test this, we sought to characterize its role in pathogenesis *in vivo* using an established pregnant mouse model of ascending vaginal infection and *ex vivo* with primary human PMs. Here, we demonstrate that *npx* is required for GBS survival in PMs and full virulence and invasion of gravid reproductive tissues *in vivo*. We have also demonstrated that *npx* is required to induce specific inflammatory cytokine profiles expressed during GBS infection.

## Materials and Methods

### Bacterial strains and culture conditions

*S. agalactiae* strain GB00112 (GB112), which represents the wild-type (WT) or parental strain, was utilized in this study. GB112 is a clinical, sequence type (ST)-12, capsular (cps) serotype III strain isolated from a rectovaginal swab of a post-partum patient (40). We have previously examined GB112 for interaction with host macrophages and fetal membranes (39, 41). Isogenic GB112 mutants were constructed previously. These include a *npx* deletion mutant (Δ*npx*) and a complemented Δ*npx* mutant harboring a plasmid containing the *npx* locus (Δ*npx:C*), was previously described (39). Bacterial strains were grown on tryptic soy agar plates supplemented with 5% sheep blood or in Todd-Hewitt broth (THB) at 37°C. Derivatives harboring the pLZ12 plasmid were grown in media supplemented with 3 µg/mL chloramphenicol. *E. coli* DH5α strains used for the mutation and complementation process were grown in LB broth or agar supplemented with either 150 µg/mL erythromycin or 20 µg/mL chloramphenicol when necessary.

### Purification of placental macrophages (PMs)

De-identified placental tissue was collected from non-laboring women who delivered healthy, full-term infants by Caesarian section at Vanderbilt University Medical Center with approval from the Vanderbilt University Medical Center Institutional Review Board (VUMC IRB #181998). PMs were isolated according to our previously published methods (41, 42). Briefly, villous core tissue was macerated and enzymatically digested with hyaluronidase, collagenase, and DNase (Sigma-Aldrich) before being strained through a stainless-steel filter and suspended in RPMI with 4-(2-hydroxyethyl)-1-piperazineethanesulfonic acid (HEPES), L-glutamine, and fetal bovine serum supplemented with antibiotic and antifungal factors. Cells were filtered and centrifuged, and CD14^+^ cells were isolated using the magnetic MACS Cell Separation system with CD14 microbeads (Miltenyi Biotec). Cells were incubated in RPMI 1640 medium (ThermoFisher) with 10% charcoal stripped fetal bovine serum (ThermoFisher) and 1% antibiotic/antimycotic solution (ThermoFisher) overnight at 37°C in 5% carbon dioxide. The following day, PMs were suspended in RPMI 1640 medium without antibiotic/antimycotic and distributed into polystyrene plates. Cells were seeded at a density of 200,000 cells per well in a polystyrene, 24-well culture plate in RPMI with 1 % antibiotic/antimycotic solution and 10 % charcoal dextran FBS (RPMI +/+), and then incubated for 24 hr in a humidified atmosphere at 37 °C and 5 % CO_2_.

### Co-culture of GBS with placental macrophages and survival assays

PMs were cultured in RPMI 1640 medium (ThermoFisher) with 10% charcoal stripped fetal bovine serum (ThermoFisher) and 1% antibiotic/antimycotic solution (ThermoFisher) overnight at 37°C in room air supplemented with 5% carbon dioxide. Co-cultured cells were incubated at 37°C in air supplemented with 5% carbon dioxide for 1 to 24 hr with media free of antibiotic/antimycotic solution. Macrophages were inoculated at a multiplicity of infection (MOI) of 10:1 bacteria to host cells for 1 hr. Co-cultures were washed with sterile media, resuspended in fresh medium containing 100 μg/mL of gentamicin (Sigma) to kill extracellular bacteria, and further incubated for 4 hr at 37°C. Gentamicin kills extracellularly located GBS but is limited in its ability to gain access to intracellular organisms. Subsequently, the samples were extensively washed with sterile PBS, and dislodged with trypsin (0.05% Trypsin-EDTA 1x, Gibco). Following collection of the PMs, the cells were lysed by the addition of 1 mL of dH_2_O. Cellular cytoplasmic contents were serially diluted in PBS and plated on blood agar plates to determine the number of viable intracellular bacteria. Samples containing only bacteria were used to estimate the efficacy of antibiotic killing as a control experiment (43).

### Transmission Electron Microscopy (TEM) analyses

Co-cultures of GBS and macrophages were subjected to primary fixation with 2.5% glutaraldehyde, 2.0% paraformaldehyde, in 0.05 M sodium cacodylate buffer at room temperature for 24 hr. Subsequently, samples were washed three times with 0.05 M sodium cacodylate buffer and subjected to a secondary fixation step with 0.1% osmium tetroxide for 15 min. Samples were washed three times with 0.05 M sodium cacodylate buffer before being sequentially dehydrated with increasing concentrations of ethanol. After dehydration, samples were embedded in resin, polymerized, and sectioned into 70-90 nm sections via ultramicrotomy. Sections were lifted onto nickel or copper 100 mesh grids (Electron Microscopy Sciences) and secondarily stained with 1% phosphotungstic acid. Grids were imaged with a Philips/FEI T12 transmission electron microscope to visualize intracellular bacteria. Bacteria were enumerated by blinded analysis using ImageJ software package.

### Ascending vaginal infection model

GBS infection of pregnant mice and subsequent analyses were performed as previously described (15, 44). Briefly, C57BL6/J mice were purchased from Jackson Laboratories and mated in harem breeding strategies (1 male to 3-4 females) overnight. Pregnancy was confirmed by the presence of a vaginal mucus plug establishing the embryonic date (E0.5) the following day. On embryonic day 13.5 (E13.5), pregnant dams were anesthetized via inhalation of isoflurane and vaginally infected with 5×10^2^-10^4^ colony forming units (CFU) in 0.05 mL of THB plus 10% gelatin. Uninfected controls were also maintained. Infections were allowed to progress until term (E21.5) for disease progression studies, or until embryonic day 15.5 (E15.5) for bacterial and immunological assays. Animals were euthanized by carbon dioxide asphyxiation and necropsy was performed to harvest reproductive tissues including uterus, placenta, decidua, amnion, and fetus tissues.

### Quantitative culture evaluation of bacterial burden

To determine bacterial burden in reproductive tissues, quantitative culture methods were employed as previously described (44). Briefly, reproductive tissues were weighed and placed in sterile THB. Tissues were homogenized and subjected to serial dilution and plated onto blood agar to quantify bacteria (CFU/mg) in host tissue. Plates were incubated at 37°C in air supplemented with 5% carbon dioxide overnight and individual colonies were counted to quantify CFU.

### Evaluation of bacterial growth in amniotic fluid

To determine bacterial growth in human or mouse amniotic fluid, quantitative culture methods were employed as described above. Briefly, human amniotic fluid was pipetted off of the surface of placentas from term, non-laboring c-section deliveries. Mouse amniotic fluid was collected after necropsy on embryonic day 15.5. All amniotic fluid samples were stored at -80°C until utilization for growth assays. Amniotic fluid samples (100 L) were placed into 96-well plates and a 1:100 dilution of overnight bacterial culture (either WT GBS, Δ*npx* mutant, or the complemented derivative) was inoculated into the amniotic fluid. Samples were incubated overnight (16-24 hours) at 37°C in room air supplemented with 5% CO_2_. The following day, bacterial viability was enumerated via serial dilution and plating onto bacteriological medium (blood agar plates). Plates were incubated at 37°C in room air supplemented with 5% CO_2_ overnight and the following day bacterial colonies were counted to enumerate viable bacterial cells.

### Evaluation of cytokine response to GBS infection

Mouse reproductive tissues, maternal sera, and amniotic fluid were analyzed by multiplex cytokine assays. Mouse tissues were placed in 0.5 mL of sterile PBS or THB+ 10 mg/mL penicillin and homogenized and passed through a 0.22 µm filter. Samples were frozen at -80°C or on dry ice until analyses were performed. Samples were analyzed by Eve Technologies via multiplex cytokine array (Eve Technologies, Alberta, Canada) as previously described (45). Validation of host targets for specific cytokines (IL-1β, IL-6, KC, and TNF-α) were performed by sandwich ELISA (AbCam) as indicated in our prior study (46).

### Ethics statement

This study was carried out in accordance with the recommendations of the Vanderbilt University Medical Center Institutional Review Board. This protocol was approved by the Institutional Review Board (IRB #181998 and #00005756). All animal experiments were performed in accordance with the Animal Welfare Act, U.S. federal law, and NIH guidelines. All experiments were carried out under a protocol approved by Vanderbilt University Institutional Animal Care and Use Committee (IACUC: M/14/034, M/17/002, and V2000089), a body that has been accredited by the Association of Assessment and Accreditation of Laboratory Animal Care Act (AAALAC).

### Statistical analyses

Statistical analysis of parametric data with more than two groups was performed using a one-way ANOVA with either Tukey’s or Dunnet’s *post hoc* correction for multiple comparisons; all reported *P* values were adjusted to account for multiple comparisons. For parametric data with two groups, a Student’s *t* test or one-way ANOVA were used. *P* values of ≤0.05 were considered significant. Non-parametric data (such as log-transformed CFU data) were analyzed by Mann-Whitney U or Kruskal-Wallis tests. Disease outcome data including survival curves were analyzed using the Mantel-Cox Log-rank test and Gehan-Breslow-Wilcoxon test. All data analyzed in this work were derived from at least three biological replicates (representing different human or mouse samples). Statistical analyses were performed using GraphPad Prism 9 (GraphPad Software Inc.).

### Data Availability

Data are available upon reasonable request to the authors.

## Results

### Transmission electron microscopy visualization of intracellular GBS in placental macrophages

TEM was used to visualize intracellular GB112 and isogenic mutants in PMs. This analysis revealed that fewer intracellular Δ*npx* cells were observed compared to WT GB112 and the isogenic Δ*npx:c* complemented derivative **(Figure 1A)**. Indeed, PM samples infected with WT GB112 averaged 17 bacterial cells per macrophage, while PMs infected with Δ*npx* averaged 5 bacterial cells per macrophage. This result was reversed via genetic complementation of the *npx* allele *in trans,* which infected an average of 16 bacterial cells per macrophage (**Figure 1B**). Quantitative culture analyses of living bacterial cells in PMs via gentamicin protection assays demonstrated that an average of 1.1×10^7^ WT GBS survived per 5×10^5^-1×10^6^ macrophages, compared to an average of 2.0×10^5^ for Δ*npx* (P < 0.05, one-way ANOVA *post hoc* Tukey’s test) **(Figure 1C)**. Meanwhile, complementation with the Δ*npx:c* vector restored survival inside macrophages, with an average of 1.4×10^7^ cells surviving per 5×10^5^-1×10^6^ macrophages, a result which was comparable to WT GB112 (P<0.01, one-way ANOVA *post hoc* Tukey’s test). Taken together, *npx* is plays in important role in persistence and survival within PMs.

**Figure 1.**
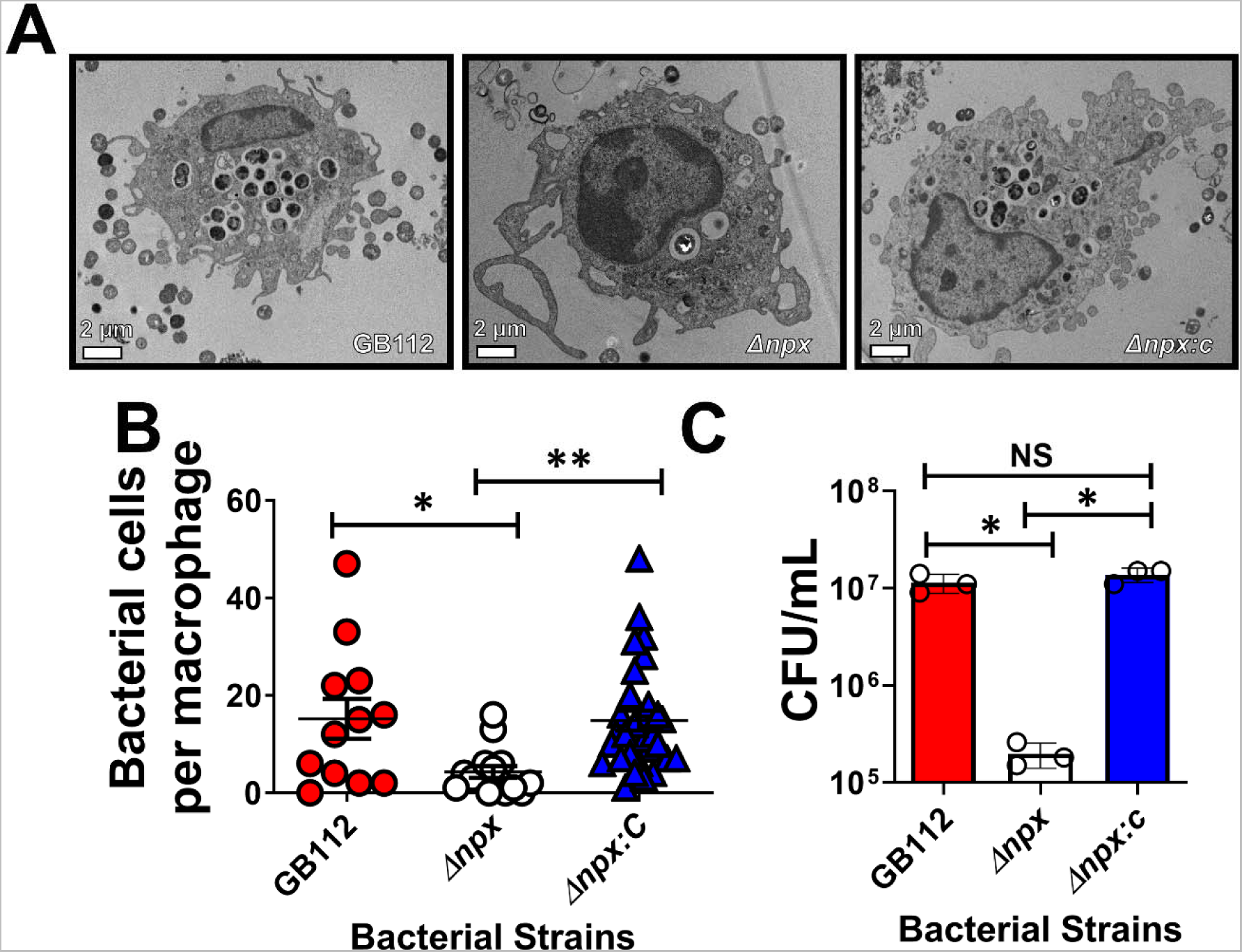
Analysis of GBS survival in primary human placental macrophage cells. A) Transmission electron microscopy analyses reveal that *npx* is required for GBS survival in placental macrophages. B) Enumeration of bacterial cells per placental macrophage via electron microscopy analyses indicates WT GBS (GB112, red) survives within primary human placental macrophages, but an isogenic Δ*npx* mutant (Δ*npx,* white) is attenuated in intracellular survival compared to both the parental (GB112) and complemented derivatives (Δ*npx:c,* blue). C) Quantitative culture of bacterial survival by gentamicin protection assay reveals *npx* is required for GBS survival in placental macrophages. *P<0.05, **P<0.01, one-way ANOVA with Tukey’s *post hoc* multiple comparisons test, NS= statistically indistinguishable; 3 biological replicates.

### Placental macrophage cytokine responses to intracellular GBS infection

Because we observed a difference in intracellular survival amongst GB112 (WT), Δ*npx*, and Δ*npx:c*, we hypothesized that there may be differences in cytokine production by PMs. Following overnight infection of PMs with GBS and its isogenic mutants, cell supernatants were collected, and cytokine quantifications were performed. Intracellular infection with WT GB112 resulted in increased secretion of a variety of proinflammatory cytokines, including CXCL1, IL- 1RA, IL-1α, IL-1β, IL-6, IL-8, MCP-1, MIP-1α, MIP-1B, and TNFα **(Figure 2)**. Nonetheless, these cytokines were similarly increased in PMs infected with Δ*npx* and Δ*npx:c*, suggesting that *npx* does not influence the production of proinflammatory cytokines by PMs ex *vivo*.

**Figure 2.**
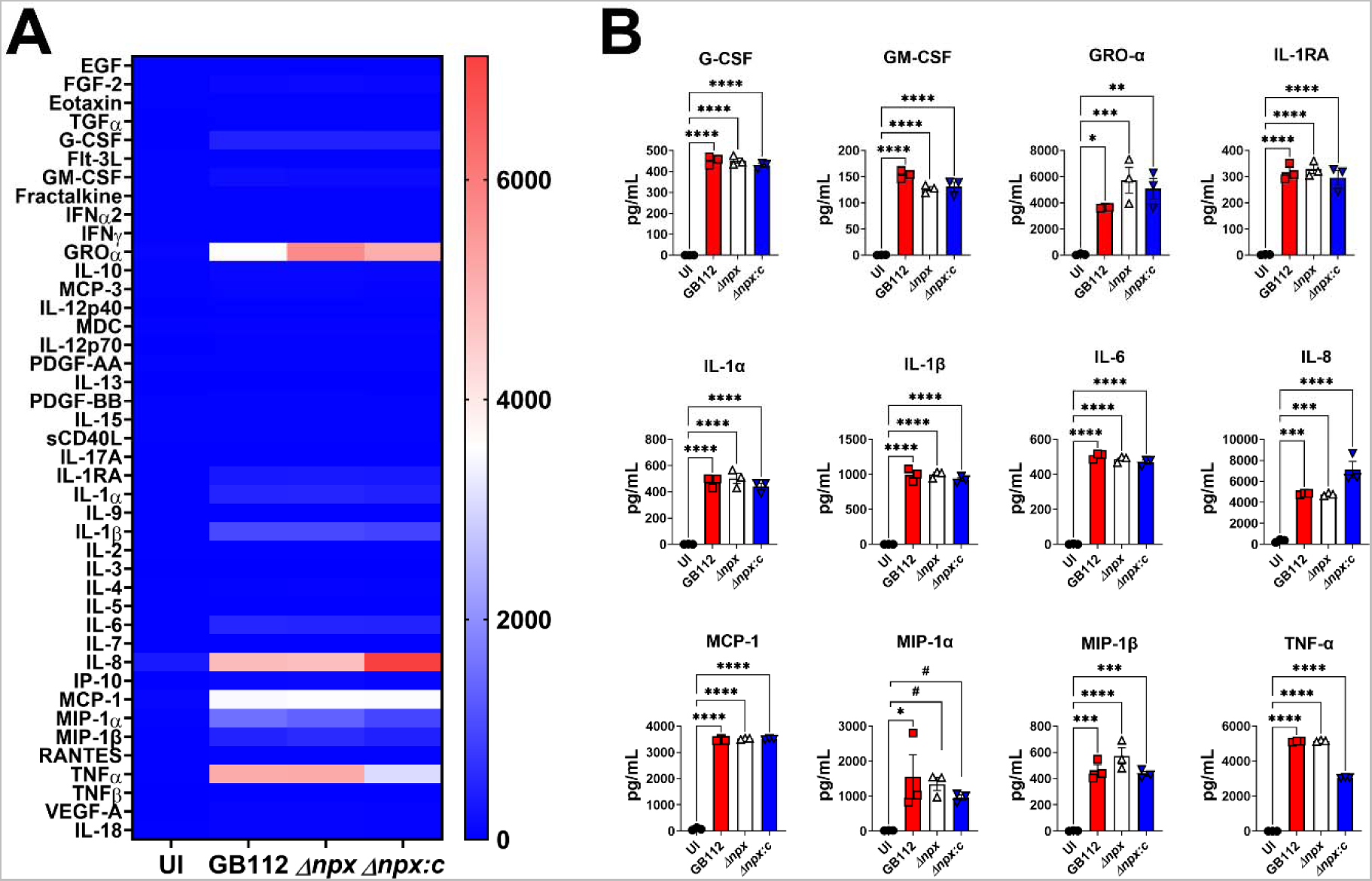
GBS *npx* expression is dispensable for initiation of proinflammatory cytokine production by human placental macrophages. A) Heat map results (blue = low levels and red = high levels) of multiplex cytokine analyses of primary human placental macrophage secreted fractions after co-culture with wild-type GB112 (GB112), Δ*npx* isogenic mutant (Δ*npx*), or the isogenic Δ*npx* complemented derivative (Δ*npx:c*) reveal GBS infection induces production of multiple proinflammatory cytokines compared to uninfected controls (UI). B) Quantitation of cytokine levels reveals significantly enhanced production of G-CSF, GM-CSF, GRO-α, IL-1RA, IL-1α, IL-1β, Il-6, IL-8, MCP-1, MIP-1α, MIP-1β, and TNF-α in macrophages co-cultured with GB112 (red), Δ*npx* (white), Δ*npx:c* (blue) compared to uninfected controls (UI, black). *P<0.05, **P<0.01, ***P<0.001, ****P<0.0001, one-way ANOVA with Tukey’s *post hoc* multiple comparisons test. #P<0.05, Student’s *t* test with Welch’s correction; 3 biological replicates.

### GBS *npx* is associated with preterm birth in a mouse model of ascending infection

To assess the role of GBS *npx in vivo* in the context of ascending pregnancy infection, we utilized a pregnant mouse model initiated by intravaginal GBS infection (44). Previous work demonstrated that in this model, ascending vaginal GBS infection results in bacterial invasion, inflammation of reproductive tissues, PPROM, preterm birth, as well as maternal and neonatal demise (44). Hence, we hypothesized that Npx could be important for GBS evasion of innate immune responses in the gravid reproductive tract. Our results (**Figure 3**) demonstrate that mice infected with WT GB112 had higher incidences of membrane rupture (PPROM) and preterm birth (PTB) than uninfected and Δ*npx*-infected animals (P=0.0024, Mantel-Cox Log-rank test and P=0.0067, Gehan-Breslow-Wilcoxon test). Notably, all GB112-infected animals experienced PPROM or PTB by 6 days post-infection. Complementation of the *npx* locus *in trans* resulted in 65% PTB and PPROM by 6 days post-infection; these results were statistically indistinguishable from cohorts infected with WT GB112 (P=0.0766, Mantel-Cox Log-rank test and P=0.1135, Gehan-Breslow-Wilcoxon test). By 7 days post-infection, GB112-infected animals had a 100% maternal mortality, a result that was significantly higher compared to uninfected controls and Δ*npx*-infected cohorts, which exhibited 0% maternal mortality by 8 days post-infection (P=0.0164, Mantel-Cox Log-rank test and P=0.0442, Gehan-Breslow-Wilcoxon test). Complementation *in trans* partially decreased the mortality rate (33% maternal death by 8 days post-infection), which was statistically indistinguishable from WT GB112 (P=0.2205, Mantel-Cox Log-rank test and P=0.4190, Gehan-Breslow-Wilcoxon test).

**Figure 3.**
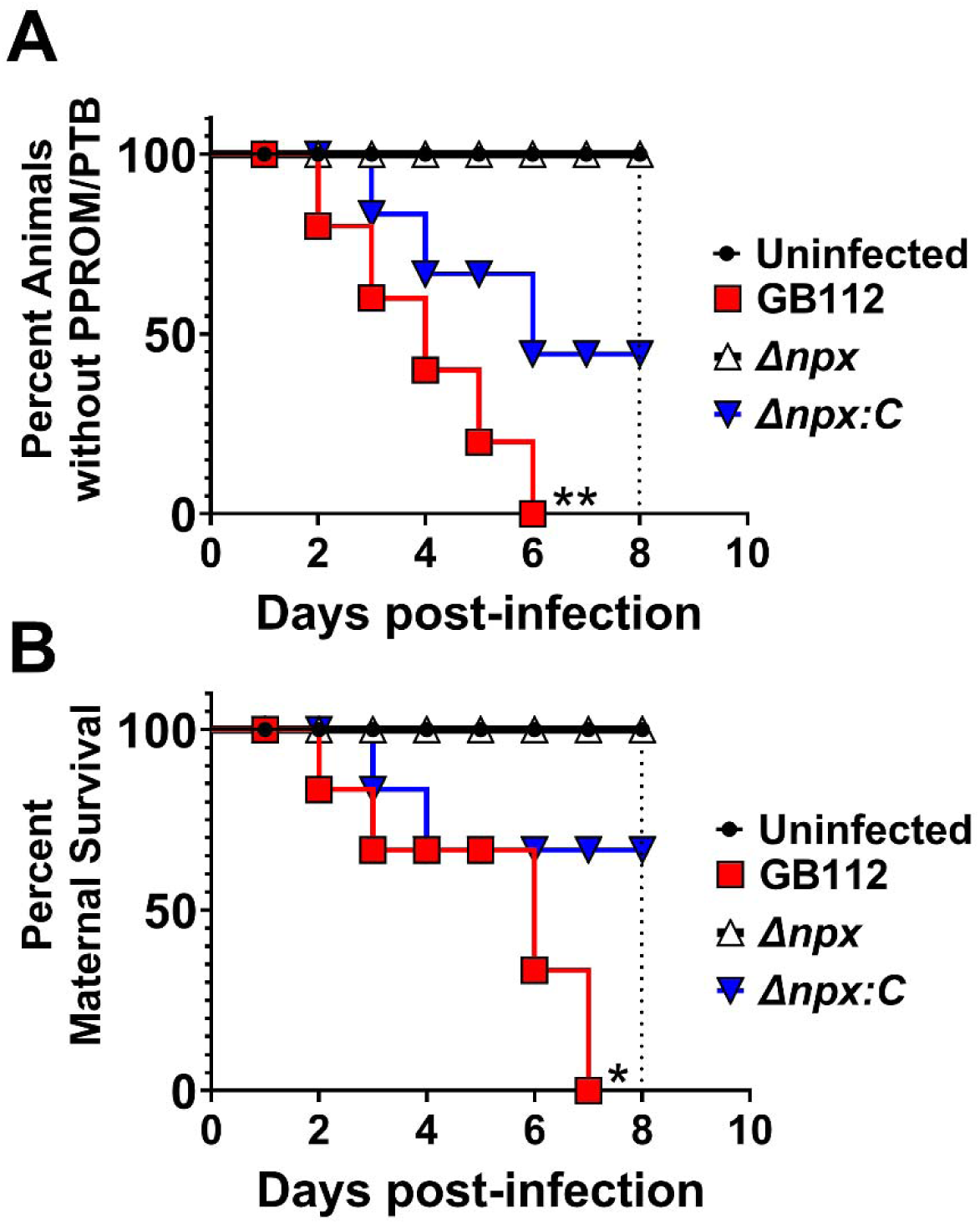
Analysis of the role of GBS *npx* in disease progression in a mouse model of ascending vaginal infection during pregnancy. A) Evaluation of the incidence of (percent of animals exhibiting) preterm premature rupture of membrane (PPROM) and preterm birth (PTB) over time (8 days post-infection and term for gestation). Wild-type GB112-infected animals (red) had higher incidences of membrane rupture (PPROM) and preterm birth (PTB) than uninfected (black) and Δ*npx*-infected animals (white), with 100% of the GB112-infected cohort experiencing PPROM or PTB by 6 days post-infection. Complementation of the *npx* locus *in trans* (Δ*npx:c,* blue) resulted in 65% PTB and PPROM by 6 days post-infection. B) Evaluation of maternal mortality over time (8 days post-infection and term for gestation) GB112-infected animals had a 100% maternal mortality by 7 days post-infection; a result that was significantly higher compared to uninfected controls and Δ*npx*-infected cohorts, which exhibited 0% maternal mortality by 8 days post-infection. Complementation *in trans* partially restored the mortality (33% maternal death by 8 days post-infection); (n=6-9 dams total from 3 separate experiments, *P<0.05, **P<0.01, Mantel-Cox Log-rank test and Gehan-Breslow-Wilcoxon test).

### Analysis of bacterial burden in reproductive tissues

To quantify the bacterial burden and classify host responses within reproductive tissues, intravaginal infections were performed using an infectious dose of 5×10^2^ to 1×10^3^ CFU. Mice were sacrificed 48 hr post-infection to collect reproductive tissues for bacteriological and immunological analyses. Bacterial burden was evaluated by quantitative culture methods for reproductive tissues including uterine, decidual, placental, amnion, and fetal tissues **(Figure 4)**. The Δ*npx* mutant exhibited a 3.5-log decrease in burden in uterine tissue compared to the parental strain (P<0.01, one-way ANOVA with Tukey’s *post hoc* multiple comparisons test). Genetic complementation *in trans* resulted in a significant increase in bacterial burden (P<0.05, one-way ANOVA with Tukey’s *post hoc* multiple comparisons test) compared to the Δ*npx* mutant. The Δ*npx* mutant exhibited a 7.5-log decrease in burden in decidual tissue compared to the parental strain (P<0.0001, one-way ANOVA with Tukey’s *post hoc* multiple comparisons test). Genetic complementation *in trans* resulted in a significant increase in bacterial burden (P<0.0001, one-way ANOVA with Tukey’s *post hoc* multiple comparisons test) compared to the Δ*npx* mutant. The Δ*npx* mutant exhibited a 7.2-log decrease in burden in placental tissue compared to the parental strain (P<0.0001, one-way ANOVA with Tukey’s *post hoc* multiple comparisons test). Genetic complementation *in trans* resulted in a significant increase in bacterial burden (P<0.0001, one-way ANOVA with Tukey’s *post hoc* multiple comparisons test) compared to the Δ*npx* mutant. The Δ*npx* mutant exhibited a 5.7-log decrease in burden in amnion tissue compared to the parental strain (P<0.0001, one-way ANOVA with Tukey’s *post hoc* multiple comparisons test). Genetic complementation *in trans* resulted in a significant increase in bacterial burden (P<0.001, one-way ANOVA with Tukey’s *post hoc* multiple comparisons test) compared to the Δ*npx* mutant. The Δ*npx* mutant exhibited a 6.5-log decrease in burden in fetal tissue compared to the parental strain (P<0.0001, one-way ANOVA with Tukey’s *post hoc* multiple comparisons test). Genetic complementation *in trans* resulted in a significant increase in bacterial burden (P<0.0001, one-way ANOVA with Tukey’s *post hoc* multiple comparisons test) compared to the Δ*npx* mutant. This *in vivo* study revealed that Npx is required for GBS to achieve full bacterial burden in reproductive tissues in an ascending model of infection during pregnancy.

**Figure 4.**
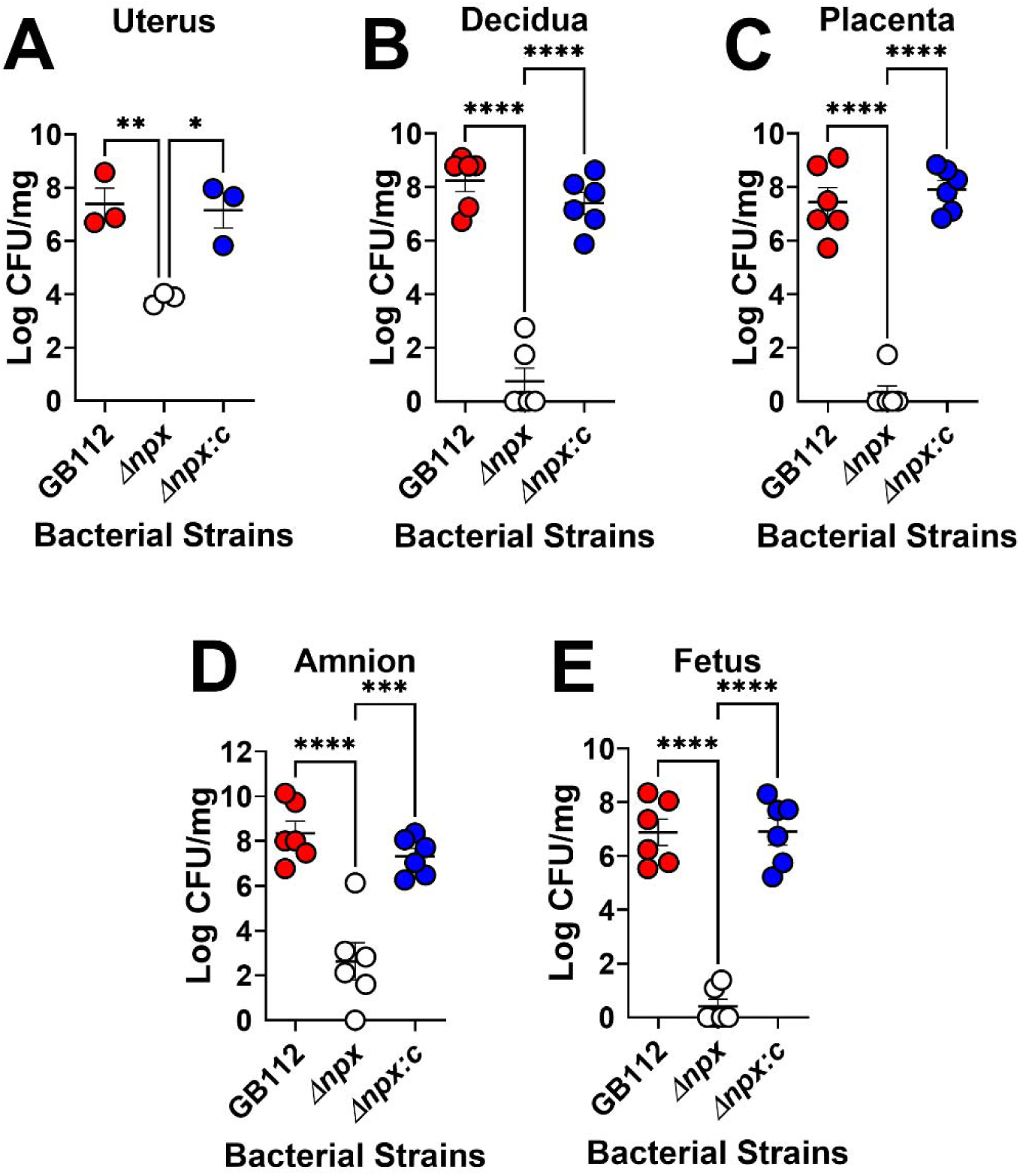
Analysis of bacterial burden within gravid reproductive tissues. Quantitative culture of bacterial burden within the A) uterus, B) decidua, C) placenta, D) amnion, and E) fetus of pregnant mice on embryonic day E15.5, two days post-infection with either WT GBS (GB112, red), an isogenic Δ*npx* mutant (Δ*npx,* white), or the isogenic complemented derivative (Δ*npx:c,* blue). GBS *npx* is required for full bacterial burden in reproductive tissues in an ascending model of infection during pregnancy. *P<0.05, **P<0.01, ***P<0.001, ****P<0.0001, one-way ANOVA with Tukey’s *post hoc* multiple comparisons test; n=3 dams total, with 1-2 fetal-placental units analyzed per dam.

### Analysis of bacterial growth in amniotic fluid

Because the Δ*npx* mutant exhibited attenuated bacterial burden within reproductive tissues compared to WT or complemented derivatives, we hypothesized that the Δ*npx* mutant might have an attenuated ability to grow in certain environments, such as amniotic fluid. To test this, we utilized human and mouse amniotic fluid and performed a 1:100 dilution inoculation with WT GBS, Δ*npx* mutant, or the complemented derivative. Samples were incubated overnight and bacterial viability was enumerated via serial dilution and quantitative culture techniques. The results indicate that the Δ*npx* mutant is attenuated in its ability to grow in human amniotic fluid and mouse amniotic fluid (P<0.001 and P<0.0001, respectively, one-way ANOVA with Tukey’s *post hoc* multiple comparisons test); results which were reversed by genetic complementation techniques (**Supplemental Figure 1**).

### Expression of GBS npx alters cytokine response to GBS infection

Because the Δ*npx* mutant showed decreased bacterial burden and disease progression in the pregnant mice, we hypothesized that this could be attributed to changes in proinflammatory cytokine production as a consequence of bacterial infection. To test this, we utilized multiplex cytokine assays to quantify the repertoire of cytokines and chemokines produced within reproductive tissues in response to GBS infection and compared them to uninfected animal tissues. Our results indicate that infection with WT GBS significantly enhanced production of IL-1β, MIP-1α, and TNF-α in the uterus, decidua, placenta, amnion, and fetus (**Figures 5-9** and **Supplemental Figures 2-6**) compared to uninfected controls (*P<0.05, one-way ANOVA, #P<0.05, Student’s *t* test). Importantly, deletion of the *npx* gene resulted in a significantly reduced production of these cytokines and chemokines compared to WT-infected samples (*P<0.05, one-way ANOVA, #P<0.05, Student’s *t* test). Similarly, G-CSF, MCP-1, MIG, and MIP-2 were significantly upregulated in the uterus, decidua, placenta, and amnion of animals infected with WT GBS compared to uninfected animals (*P<0.05, one-way ANOVA, #P<0.05, Student’s *t* test), and loss of the *npx* gene resulted in a significant reduction in production of these cytokines and chemokines compared to WT-infected samples (*P<0.05, one-way ANOVA, #P<0.05, Student’s *t* test). IL-6 and MIP-1 were upregulated in the uterus, decidua, placenta, and fetus in response to WT GBS infection compared to uninfected controls, and deletion of the *npx* gene revealed significant reduction in production of these cytokines and chemokines compared to WT-infected samples (*P<0.05, one-way ANOVA, #P<0.05, Student’s *t* test). IP-10 was upregulated in the uterus, decidua, and placenta in response to WT GBS infection compared to uninfected controls, and a significant reduction in production of these cytokines and chemokines was observed with inactivation of the *npx* gene compared to WT-infected samples (*P<0.05, one-way ANOVA, #P<0.05, Student’s *t* test). Complementation with the WT *npx* allele *in trans* restored cytokine and chemokine production similar to those observed in WT GBS-infected animals, or to levels that were significantly higher than those measured in the isogenic Δ*npx* mutant-infected samples (*P<0.05, one-way ANOVA, #P<0.05, Student’s *t* test, NS= statistically indistinguishable from WT GBS).

**Figure 5.**
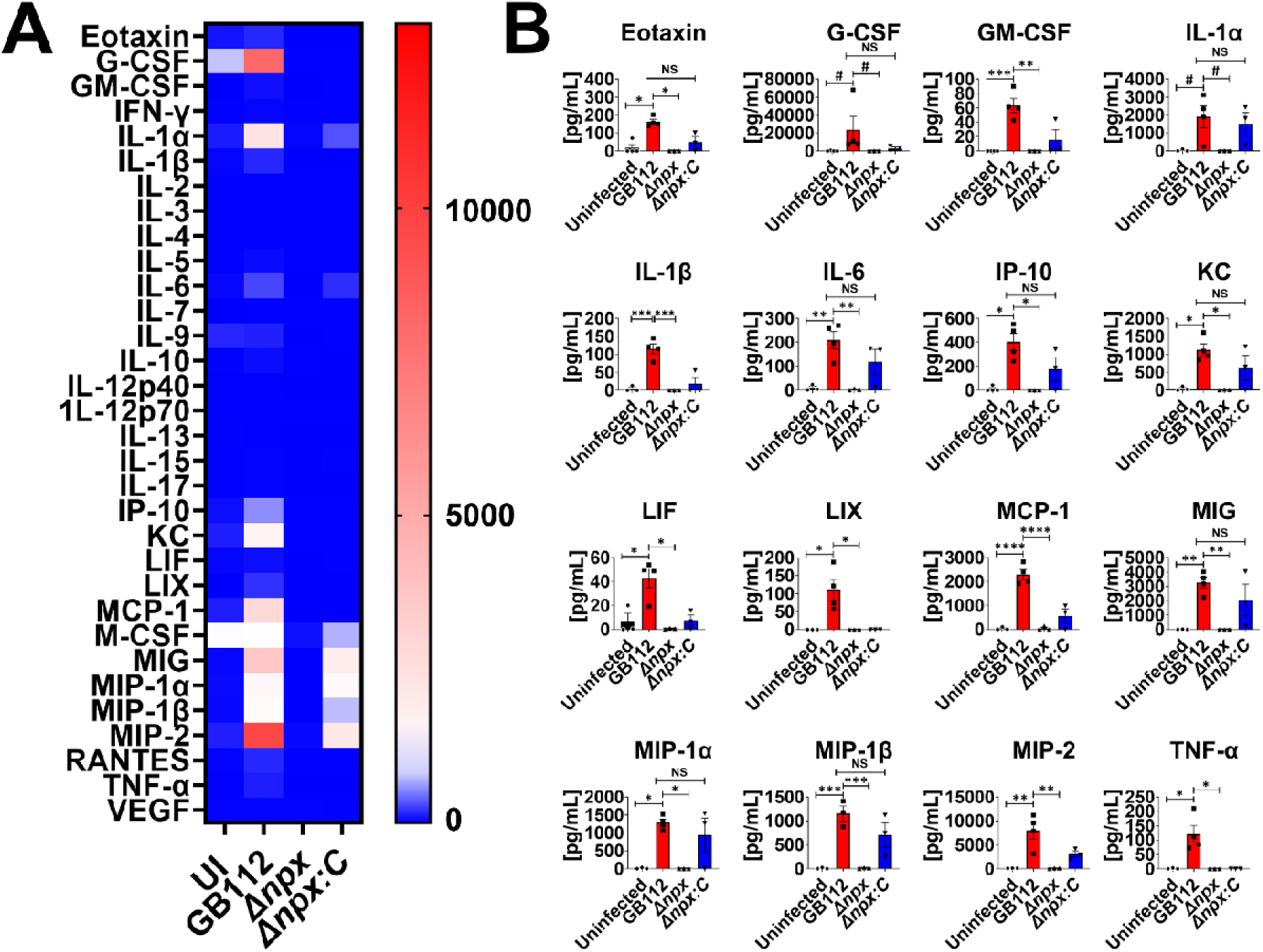
Analysis of cytokine production in uterus tissue in response to GBS infection. A) Heat map results (blue = low levels and red = high levels) of multiplex cytokine analyses of gestational tissues after ascending vaginal infection with wild-type GB112 (GB112), Δ*npx* isogenic mutant (Δ*npx*), or the isogenic Δ*npx* complemented derivative (Δ*npx:c*) as well as uninfected controls (UI). Uterine tissues were collected from pregnant mice on embryonic day E15.5, two days post- vaginal infection with GBS. B) Quantification of Eotaxin, G-CSF, GM- CSF, IL-1α, IL-1β, IL-6, IP-10, KC, LIF, LIX, MCP-1, MIG, MIP-1α, MIP-1β, MIP-2, and TNF-α levels revealed that wild-type GB112 infection (red bars) significantly enhances production of these cytokines compared to uninfected controls (black bars), however the isogenic Δ*npx* mutant (white bars) is significantly attenuated in its ability to induce these cytokines compared to the parental strain. Conversely, the complemented derivative (Δ*npx:c*, blue bars) is often statistically indistinguishable from the parental strain (NS). Bars indicate mean values +/- standard error mean with individual data points representing results from uterus tissues from individual dams. #P<0.05, Student’s *t* test, *P<0.05, **P<0.01, ***P<0.001, ****P<0.0001, one-way ANOVA with Tukey’s *post hoc* multiple comparisons test, NS= not statistically significant. Results indicate that GBS *npx* is required for full initiation of proinflammatory cytokine response in uterine tissues in an ascending model of infection during pregnancy.

**Figure 6.**
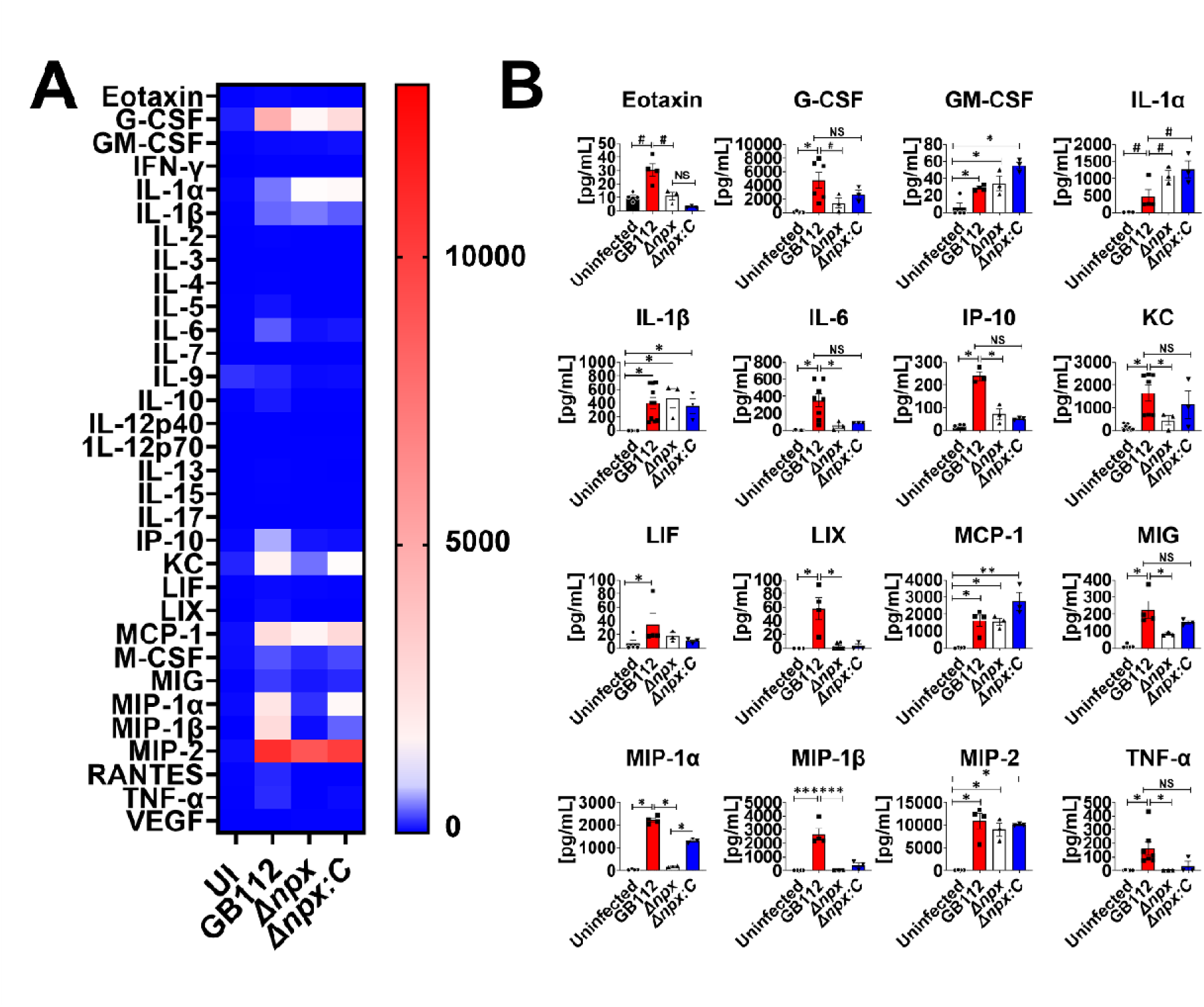
Analysis of cytokine production in decidua tissue in response to GBS infection. A) Heat map results (blue = low levels and red = high levels) of multiplex cytokine analyses of decidua tissues after ascending vaginal infection with wild-type GB112 (GB112), Δ*npx* isogenic mutant (Δ*npx*), or the isogenic Δ*npx* complemented derivative (Δ*npx:c*) as well as uninfected controls (UI). Decidua tissues were collected from pregnant mice on embryonic day E15.5, two days post- vaginal infection with GBS. B) Quantification of Eotaxin, G-CSF, GM-CSF, IL-1α, IL- 1β, IL-6, IP-10, KC, LIF, LIX, MCP-1, MIG, MIP-1α, MIP-1β, MIP-2, and TNF-α levels revealed that wild-type GB112 infection (red bars) significantly enhances production of these cytokines compared to uninfected controls (black bars), however the isogenic Δ*npx* mutant (white bars) is significantly attenuated in its ability to induce these cytokines compared to the parental strain. Conversely, the complemented derivative (Δ*npx:c*, blue bars) is often statistically indistinguishable from the parental strain (NS). Bars indicate mean values +/- standard error mean with individual data points representing results from decidual tissues from individual fetal- placental units from separate dams. #P<0.05, Student’s *t* test, *P<0.05, **P<0.01, ***P<0.001, ****P<0.0001, one-way ANOVA with Tukey’s *post hoc* multiple comparisons test, NS= not statistically significant. Results indicate that GBS *npx* is required for full initiation of proinflammatory cytokine response in decidual tissues in an ascending model of infection during pregnancy.

Infection with WT GBS resulted in significantly enhanced production of select cytokines and chemokines including: eotaxin, G-CSF, GM-CSF, IL-1α, IL-1β, IL-6, IL-15, IP-10, KC, LIF, LIX, MCP-1, MIG, MIP-1α, MIP-1β, MIP-2, and TNF-α in the uterus (**Figure 5** and **Supplemental Figure 2**), eotaxin, G-CSF, GM-CSF, M-CSF, IL-1α, IL-1β, IL-6, IL-15, IP-10, KC, LIF, LIX, MCP-1, MIG, MIP-1α, MIP-1β, MIP-2, and TNF-α in the decidua (**Figure 6** and **Supplemental Figure 3**), G-CSF, IL-1β, IL-6, IP-10, KC, MCP-1, MIG, MIP-1α, MIP-1β, MIP-2, M-CSF, and TNF-α in the placenta (**Figure 7** and **Supplemental Figure 4**), G-CSF, M-CSF, IL- 1β, KC, MCP-1, MIG, MIP-1α, MIP-2, and TNF-α in the amnion (**Figure 8** and **Supplemental Figure 5**), eotaxin, IL-1β, IL-6, KC, MIP-1α, MIP-1β, and TNF-α in the fetus (**Figure 9** and **Supplemental Figure 6**), compared to uninfected controls (*P<0.05, one-way ANOVA, #P<0.05, Student’s *t* test). Inactivation of the *npx* gene, however, evoked a significant reduction in production of many of these cytokines and chemokines compared to WT-infected samples (P<0.05, one-way ANOVA, #P<0.05, Student’s *t* test).

**Figure 7.**
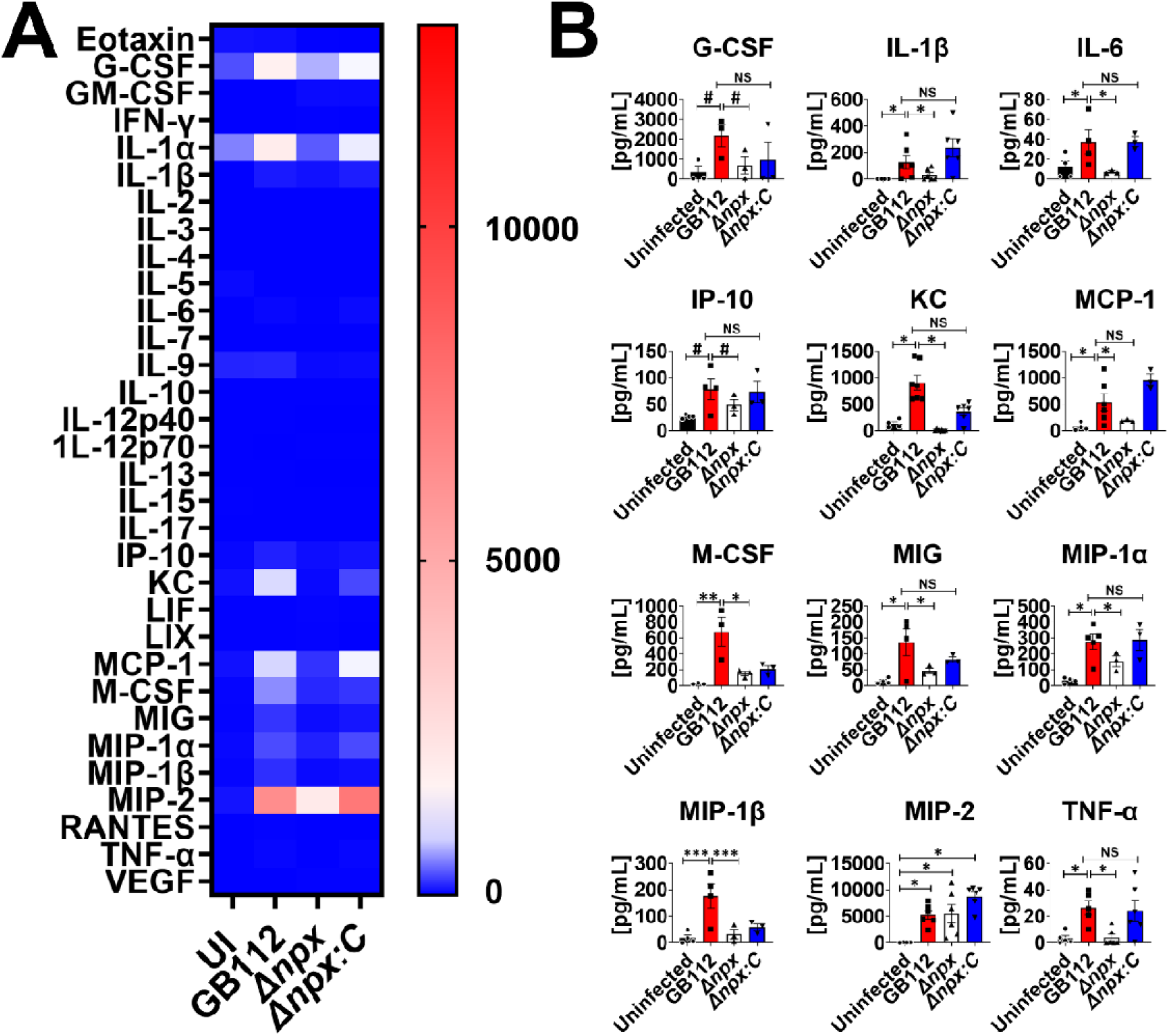
Analysis of cytokine production in placenta tissue in response to GBS infection. A) Heat map results (blue = low levels and red = high levels) of multiplex cytokine analyses of placenta tissues after ascending vaginal infection with wild-type GB112 (GB112), Δ*npx* isogenic mutant (Δ*npx*), or the isogenic Δ*npx* complemented derivative (Δ*npx:c*) as well as uninfected controls (UI). Placenta tissues were collected from pregnant mice on embryonic day E15.5, two days post- vaginal infection with GBS. B) Quantification of G-CSF, IL-1β, IL-6, IP-10, KC, MCP-1, M-CSF, MIG, MIP-1α, MIP-1β, MIP-2, and TNF-α levels revealed that wild-type GB112 infection (red bars) significantly enhances production of these cytokines compared to uninfected controls (black bars), however the isogenic Δ*npx* mutant (white bars) is significantly attenuated in its ability to induce these cytokines compared to the parental strain except MIP-2. Conversely, the complemented derivative (Δ*npx:c*, blue bars) is often statistically indistinguishable from the parental strain (NS). Bars indicate mean values +/- standard error mean with individual data points representing results from placental tissues from individual fetal-placental units from separate dams. #P<0.05, Student’s *t* test, *P<0.05, **P<0.01, ***P<0.001, ****P<0.0001, one-way ANOVA with Tukey’s *post hoc* multiple comparisons test, NS= not statistically significant. Results indicate that GBS *npx* is required for initiation of numerous proinflammatory cytokine responses in placental tissues in an ascending model of infection during pregnancy.

**Figure 8.**
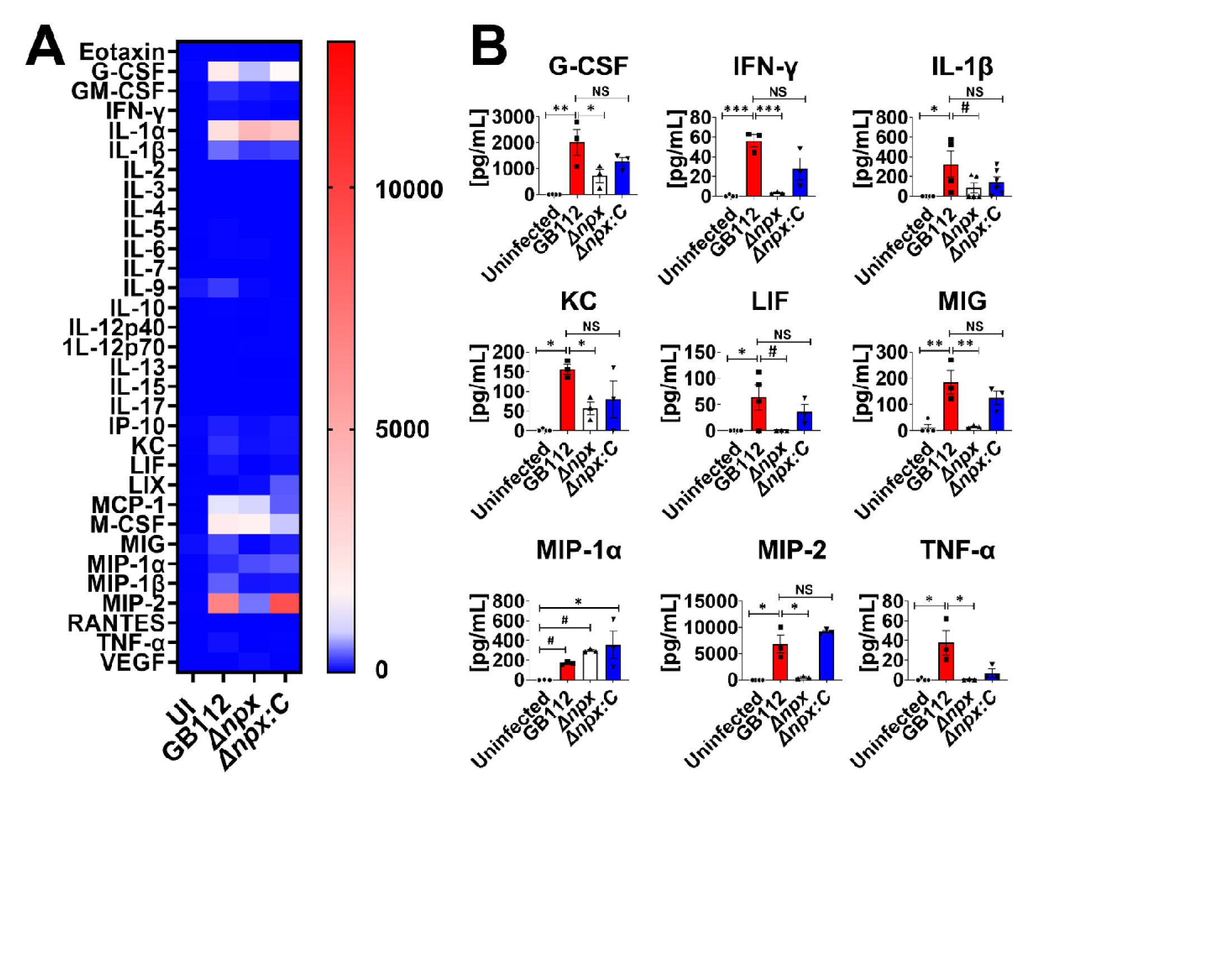
Analysis of cytokine production in amnion tissue in response to GBS infection. A) Heat map results (blue = low levels and red = high levels) of multiplex cytokine analyses of amnion tissues after ascending vaginal infection with wild-type GB112 (GB112), Δ*npx* isogenic mutant (Δ*npx*), or the isogenic Δ*npx* complemented derivative (Δ*npx:c*) as well as uninfected controls (UI). Amnion tissues were collected from pregnant mice on embryonic day E15.5, two days post- vaginal infection with GBS. B) Quantification of G-CSF, IFN-γ, IL-1β, KC, LIF, MIG, MIP-1α, MIP-2, and TNF-α levels revealed that wild-type GB112 infection (red bars) significantly enhances production of these cytokines compared to uninfected controls (black bars), however the isogenic Δ*npx* mutant (white bars) is significantly attenuated in its ability to induce these cytokines compared to the parental strain except MIP-1α. Conversely, the complemented derivative (Δ*npx:c*, blue bars) is often statistically indistinguishable from the parental strain (NS). Bars indicate mean values +/- standard error mean with individual data points representing results from amnion tissues from individual fetal-placental units from separate dams. #P<0.05, Student’s *t* test, *P<0.05, **P<0.01, ***P<0.001, ****P<0.0001, one-way ANOVA with Tukey’s *post hoc* multiple comparisons test, NS= not statistically significant. Results indicate that GBS *npx* is required for initiation of numerous proinflammatory cytokine responses in amnion tissues in an ascending model of infection during pregnancy.

**Figure 9.**
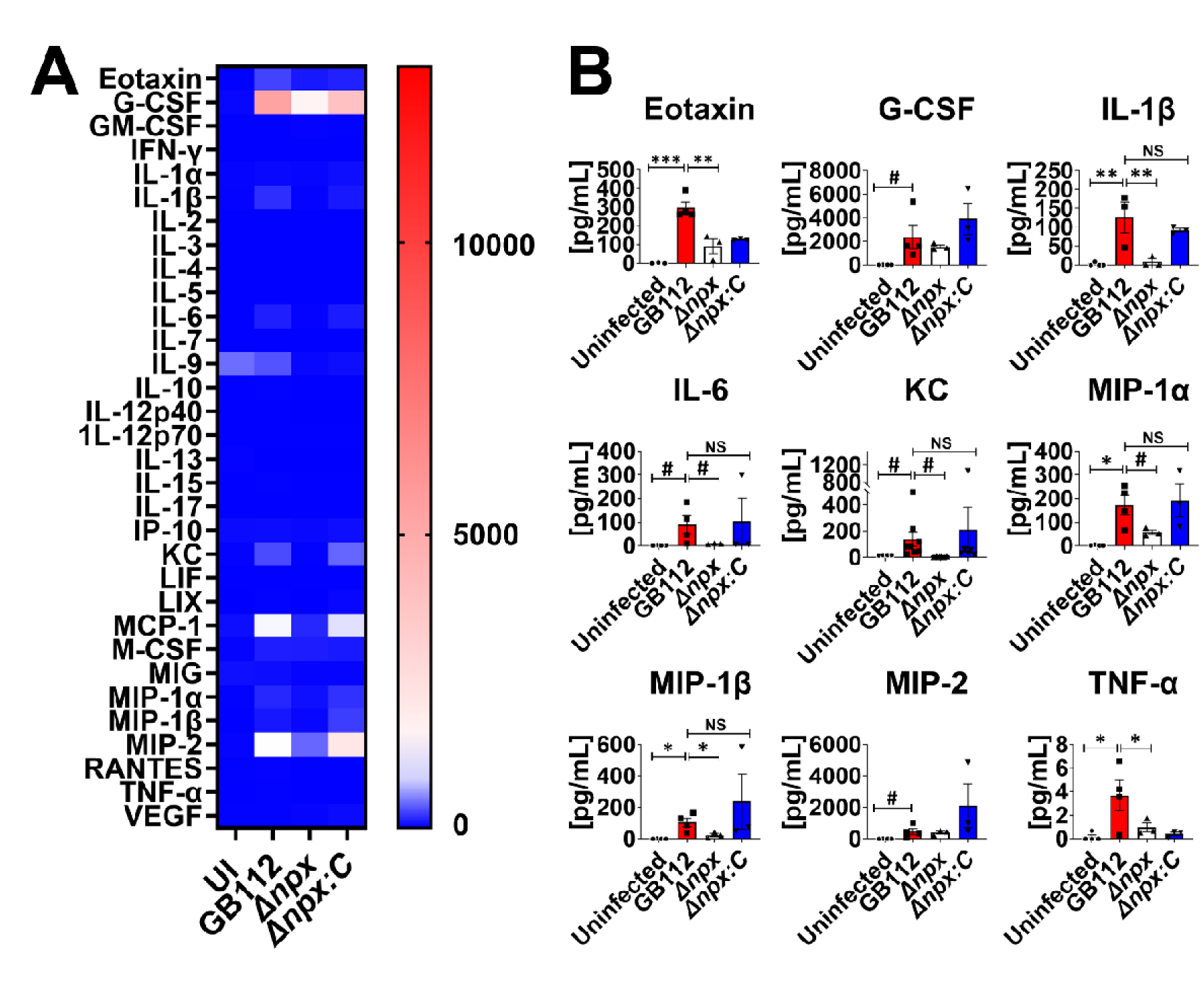
Analysis of cytokine production in fetus tissue in response to GBS infection. A) Heat map results (blue = low levels and red = high levels) of multiplex cytokine analyses of fetal tissues after ascending vaginal infection with wild-type GB112 (GB112), Δ*npx* isogenic mutant (Δ*npx*), or the isogenic Δ*npx* complemented derivative (Δ*npx:c*) as well as uninfected controls (UI). Fetus tissues were collected from pregnant mice on embryonic day E15.5, two days post-vaginal infection with GBS. B) Quantification of Eotaxin, G-CSF, IL-1β, IL-6, KC, MIP-1α, MIP- 1β, MIP-2, and TNF-α levels revealed that wild-type GB112 infection (red bars) significantly enhances production of these cytokines compared to uninfected controls (black bars), however the isogenic Δ*npx* mutant (white bars) is significantly attenuated in its ability to induce Eotaxin, IL-1β, IL-6, KC, MIP-1α, MIP-1β, and TNF-α l compared to the parental strain. Conversely, the complemented derivative (Δ*npx:c*, blue bars) is statistically indistinguishable from the parental strain (NS) in its ability to induce IL-1β, IL-6, KC, MIP-1α, and MIP-1β. Bars indicate mean values +/- standard error mean with individual data points representing results from fetal tissues from individual fetuses from separate dams. #P<0.05, Student’s *t* test, *P<0.05, **P<0.01, ***P<0.001, ****P<0.0001, one-way ANOVA with Tukey’s *post hoc* multiple comparisons test, NS= not statistically significant. Results indicate that GBS *npx* is required for initiation of numerous proinflammatory cytokine responses in fetal tissues in an ascending model of infection during pregnancy.

## Discussion

While GBS has been identified as a perinatal pathogen since the 1930s, there are still major gaps in the knowledge of the pathophysiology of infection and disease outcomes by this bacterium. We have previously observed that GBS interacts with gestational tissue macrophages, and that GBS can invade the reproductive tract in a mouse model of ascending vaginal infection during pregnancy (17, 29, 47). We sought to understand the importance of individual GBS virulence factors that influence the outcome of GBS-macrophage interactions and might also have significance in clinical outcomes during pregnancy. We previously identified that a peroxide-detoxifying enzyme NADH peroxidase, *npx,* was upregulated when GBS was cultured with THP-1 macrophages (39). We furthered expanded that work in our current study by demonstrating that GBS *npx* aids in GBS survival within primary human placental macrophages.

Macrophages represent the second most common leukocyte within fetal membrane tissues, and these cells perform many roles including regulating tissue remodeling during development and modulating maternal-fetal tolerance (48). Less is understood of the roles that macrophages may play during infection, as these cells are typically thought to be polarized to an anti-inflammatory M2 tolerogenic state (28). Some recent studies have noted that in response to bacteria, these cells change their polarization towards a more inflammatory M1 phenotype (49). GBS has evolved mechanisms such as a capsule to evade phagocytosis by placental macrophages (44), but once engulfed by innate immune cells, GBS deploy enhanced expression of the *npx* locus as a strategy to survive the peroxide stress encountered within the phagosome of THP-1 macrophages (39).

Macrophages are implicated as a replicative niche for a variety of bacteria including *Pseudomonas aeruginosa* (50), *Yersinia pestis* (51), *Brucella neotomae* (52), *Escherichia coli* (53), *Neisseria gonorrhoeae* (54), and *Legionella pneumophila* (55). Recent work has demonstrated that *S. pneumoniae* can survive and replicate within splenic macrophages, which serve as a reservoir for septicemia (56), and that Group A *Streptococcus* can survive and replicate within human macrophages (57). These results mirror what we observed with GBS in primary human PMs.

Furthermore, macrophages have been implicated as a potential Trojan Horse aiding in dissemination of a variety of microbial pathogens including *Candida albicans* (58), *Mycobacterium tuberculosis* (59), *Toxoplasma gondii* (60), *Staphylococcus aureus* (61), *Cryptococcus neoformans* (62), and *Chlamydia trachomatis* (63). Depletion of host macrophages impedes *Chlamydia* and GBS dissemination in the reproductive tract (47, 64), demonstrating the important role that intracellular bacterial survival within macrophages plays in bacterial invasion of the reproductive tract. Our previous results indicate that GBS utilizes *cadD*, a metal resistance determinant, to circumnavigate metal stress within placental macrophages, and enhance bacterial ascension and invasion of the gravid reproductive tract (47). Similarly, our current work demonstrates that GBS utilizes *npx*, a peroxide resistance determinant, to circumnavigate peroxide stress within placental macrophages, and to aid in ascending infection and disease progression during pregnancy. Taken together, these results support the hypothesis that GBS could exploit macrophages as a Trojan Horse to aid in promoting invasive bacterial infections during pregnancy.

Upon recognition of pathogens by immune pattern recognition receptors (PRR), such as toll-like receptors, a cascade of responses by the macrophage are initiated. One mechanism of defense is phagocytosis of the bacterial cell and chemical assault within the phagosome via deployment of peroxides and reactive oxygen species (65). Highly reactive oxygen species can damage macromolecules including lipids, proteins, and nucleic acids, ultimately leading to cell death (33). Within the reproductive tract, invading pathogens, including *C. trachomatis* (66) and *N. gonorrhoeae* (67), are assaulted with reactive oxygen species. There is an association between *Chlamydia* and spontaneous abortion, with oxidative stress being implicated, highlighting the oxidative response in the reproductive tract against invading pathogens (66). Additionally, the presence of ROS has been demonstrated in human amniotic fluid collected in the second and third trimester of gestation, as well (68). This likely presents an environmental challenge for bacteria which are highly sensitive to oxidative stress; results that are supported by the survival and growth defect observed in the Δ*npx* mutant compared to the wild-type and complemented derivatives grown in human or amniotic fluid.

In response to this, bacterial pathogens have evolved a range of mechanisms to overcome ROS stress inside macrophages. For instance, *S. aureus* (69), *P. aeruginosa* (70, 71), *Klebsiella pneumoniae* (72), and *M. tuberculosis* (73) all express catalase to resist oxidative killing by macrophages. Other bacterial pathogens such as *E. coli* (74), *Salmonella typhi* (75), and *Burkholderia pseudomallei* (76) express superoxide dismutase for the same purpose. GBS is catalase negative but expresses superoxide dismutase (SodA) (36). Our study demonstrates that the full repertoire of antioxidant defenses is required for invasive infection of the gravid reproductive tract, and that the *npx-*encoded NADH peroxidase aids in GBS survival within reproductive tissue macrophages and full virulence in a pregnant animal model. Interestingly, a dye-neutralizing peroxidase (DyP) has been identified in *M. tuberculosis* that is critical for bacterial survival within host macrophages as well (77). Similarly, in *Listeria monocytogenes*, peroxidases encoded within the *fri* and *ahpA* loci were each required for *L. monocytogenes* to survive acute peroxide stress, and the *fri* locus was essential for cytosolic growth within host macrophages (78). These studies further underscore the critical role that bacterial peroxidases play in the host-pathogen dialogue, specifically with respect to intracellular survival.

In addition to aiding GBS intracellular survival within host immune cells, the *npx* locus is critical for virulence *in vivo*. In our mouse model of ascending vaginal infection during pregnancy, we observed that mutants lacking *npx* showed impaired invasion into gestational tissues and cognate inflammatory responses and disease compared to the parental strain or complemented mutant, demonstrating the importance of *npx* for pathogenesis (**Figure 10**). We observed drastic reductions in bacterial burden in the uterus, decidua, placenta, amnion, and fetal tissue compartments derived from animals infected with the Δ*npx* mutant compared to the parental or complemented isogenic derivatives. These reductions in burden correlated to decreased production of proinflammatory cytokines such as IL-1β, MIP-1α, and TNF-α in all the examined tissue compartments of the gravid reproductive tract. Interestingly, diminution of proinflammatory cytokine production and bacterial burden was associated with cognate decreases in adverse pregnancy outcomes such as PPROM, preterm birth, and maternal demise. Previous work has linked high levels of IL-1β, MIP-1α, and TNF-α with enhanced risk for preterm birth (79). It is likely that expression of proinflammatory cytokines perturbs maternal tolerance of the semi-allogenic fetus, leading to enhanced risk of adverse pregnancy outcomes (80). Recently, interest has piqued in exploiting the NLRP3 inflammasome pathway as a potential chemotherapeutic strategy to ameliorate risk associated with perinatal disease outcomes (81). Because expression of the *npx* locus is critical for GBS ascension of the reproductive tract and initiation of these signaling pathways which promote inflammation, a dual- targeted approach of inhibition of the GBS NADH peroxidase and the NLRP3 inflammasome pathway could prove useful in combatting GBS perinatal infections.

**Figure 10.**
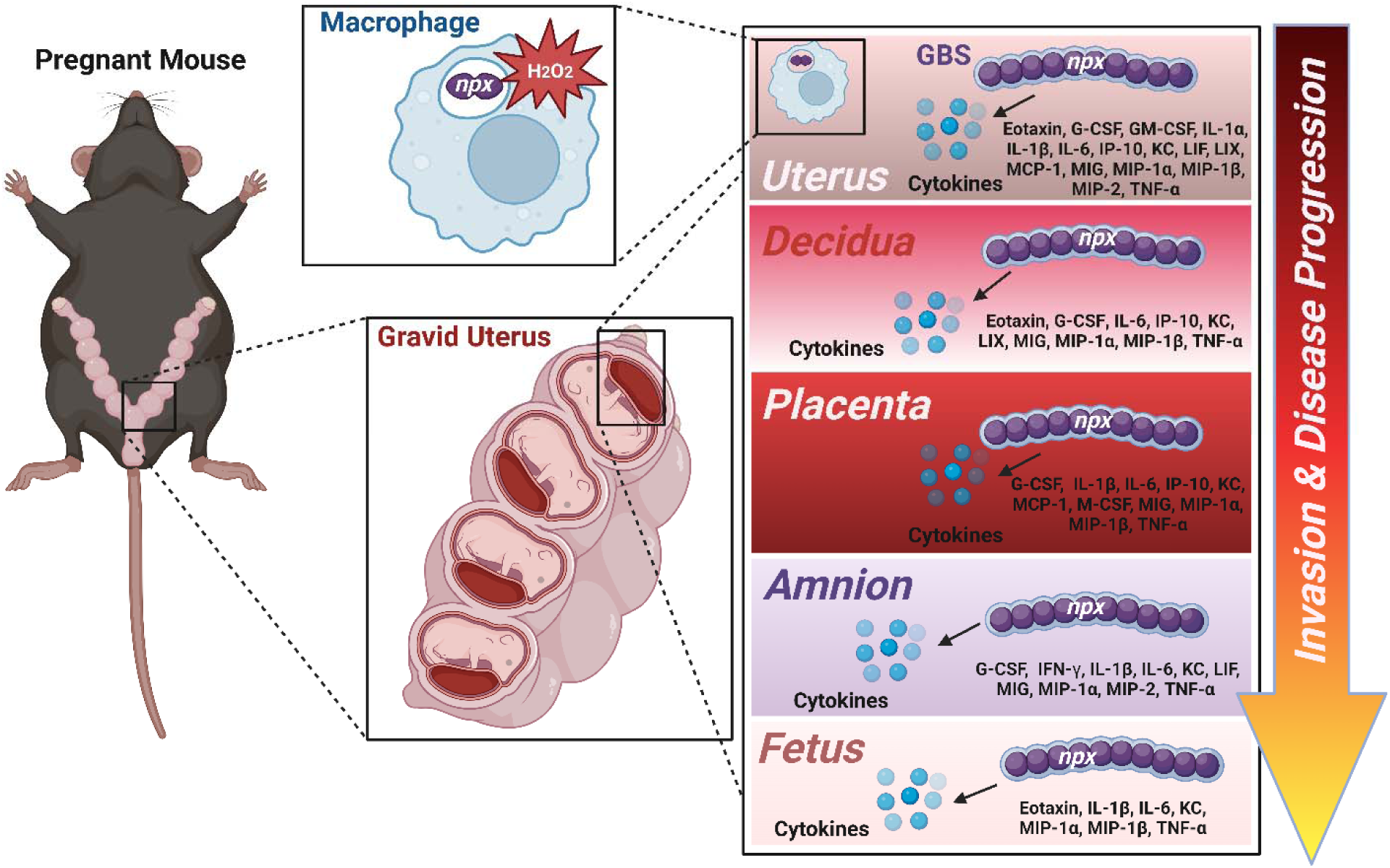
Conceptual model of *npx*-dependent invasion and proinflammatory signal initiation in ascending GBS infections during pregnancy. GBS ascends the reproductive tract by surviving within placental macrophages, in part, by encoding a NADH peroxidase to aid in detoxifying oxidative stress (such as peroxides; H_2_O_2_). GBS invasion and full burden within the gravid reproductive tract (uterus, decidua, placenta, amnion, and fetus) requires *npx* expression and triggers production of tissue compartment-specific proinflammatory cytokines resulting in inflammation and disease progression including rupture of membranes, preterm birth, and maternal demise. Image created with BioRender.com.

Intracellular infection of PMs resulted in the production of proinflammatory cytokines such as G-CSF, GM-CSF, GRO-α, IL-1RA, IL-1α, IL-1β, IL-6, IL-8, MCP-1, MIP-1α, MIP-1β, and TNF-α. However, *npx* was dispensable for this induction. This was a surprising result, considering the significant differences in bacterial load within these macrophages, underscoring that even low levels of GBS intracellular infection are sufficient to induce production of these cytokines. Production of proinflammatory cytokines by macrophages instigates the recruitment of neutrophils in mice upon GBS infection (15, 30). A similar response has also been observed in humans (30). Neutrophils aid in the clearance of bacteria by phagocytosis and subsequent killing in internal vacuoles called phagosomes (31). Finally, neutrophils excrete neutrophil extracellular traps (NETs) loaded with varying antimicrobial peptides (32). These extracellular DNA traps have been exhibited in response to GBS infection (15). Collectively, placental macrophages and neutrophils predominate the innate immune response against GBS infection.

Our studies indicate that NADH peroxidase plays an important role for the full virulence of GBS in a mouse model of ascending vaginal infection during pregnancy. Other reactive oxygen species detoxifying enzymes such as catalase and superoxide dismutase are also important for virulence for many bacterial pathogens, especially those that survive and establish replicative niches within macrophages (82, 83). Introduction of bacterial-specific inhibitors of oxidative stress response pathways may target bacteria without impacting the host and provide exciting new avenues of drug development. As such, developing small molecules or other chemotherapeutic strategies to inhibit these enzymes may be viable option to defend against these bacterial infections. Identification of these bacteria-specific inhibitors are being researched currently and include small molecules which inhibit *M. tuberculosis* catalase (84). A plausible future direction of our work may include screening pre-existing banks of small molecules against purified GBS NADH peroxidase protein or creating a crystal structure to identify regions for targeted drug design.

## Acknowledgments

This work was funded by National Institutes of Health grant NICHD R01 HD090061 (to J.A.G.) and by NIH T32 HL007411-36S1 (supporting J.L.), 2T32AI112541-06 (supporting J.D.F.), K08AI151100 (supporting R.S.D.), K12HD087023 (supporting K.N.N), and a Merit Review Award I01 BX005352-01 (to J.A.G) from the Office of Medical Research, Department of Veterans Affairs, and NSF 1847804 (to S.D.T.). Additional supported was provided by a Vanderbilt Faculty Research Scholars Award (to R.S.D), the Global Alliance to Prevent Prematurity and Stillbirth project N015615 (to D.M.A. and S.D.M.), NIH R01AI134036 (to D.M.A. and J.A.G.), and the March of Dimes (to D.M.A.). Core Services were performed through both Vanderbilt University Medical Center’s Digestive Disease Research Center supported by NIH grant P30DK058404 Core Scholarship and Vanderbilt Institute for Clinical and Translational Research program supported by the National Center for Research Resources, Grant UL1 RR024975-01, and the National Center for Advancing Translational Sciences, Grant 2 UL1 TR000445-06. Imaging experiments were performed, in part, with the Vanderbilt Cell Imaging Shared Resource (supported by NIH grants CA68485, DK20593, DK58404, DK59637 and EY08126). The content is solely the responsibility of the authors and does not necessarily represent the official views of the NIH or any of the other supporters.

## Author Contributions

All authors listed contributed substantially to this work. Experiments were conducted, data were analyzed, and figures were prepared by J.L., R.S.D., M.A.G., R.E.M., S.K.S., J.D.F., J.A.T., L.M.R, K.N.N., M.L.K., and J.A.G. S.D.T., J.A.G., D.M.A, and S.D.M. designed the experiments and supervised these studies. J.L., R.S.D., M.A.G., R.E.M., S.K.S., S.D.T., D.M.A., S.D.M., and J.A.G. wrote the manuscript which was edited and approved by all authors prior to submission.

## Declarations of Competing Interests

The authors declare no competing interests

## Supplemental Figures

**Supplemental Figure 1.**
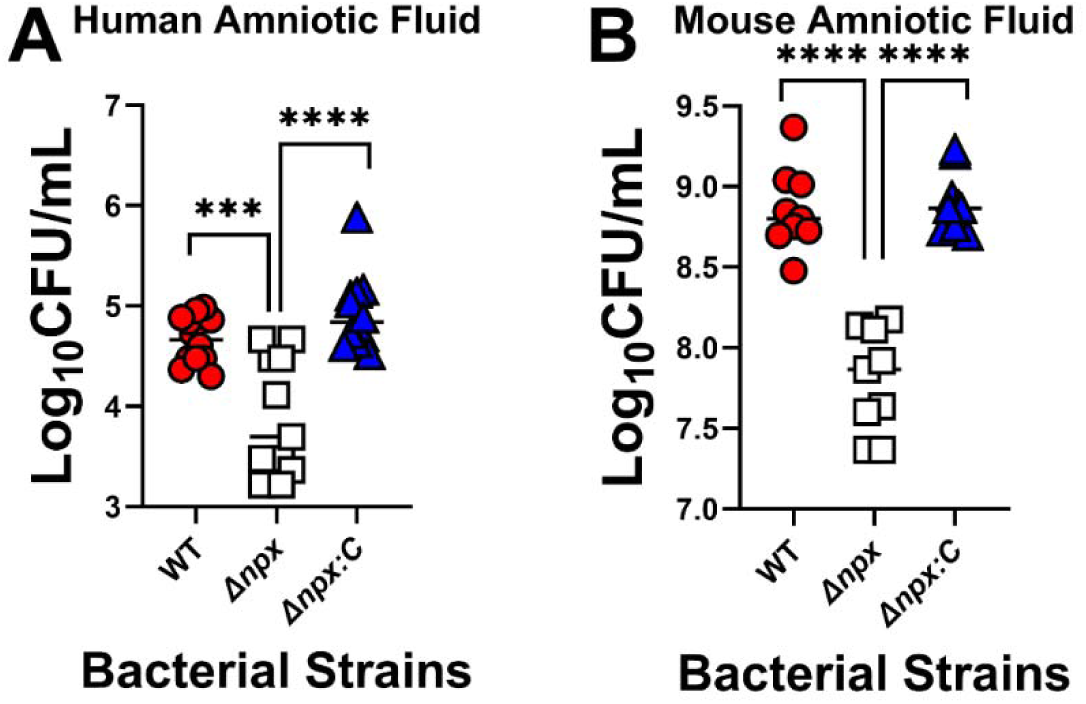
Analysis of bacterial viability in human or mouse amniotic fluid. Wild- type GB112 (WT, red circles), Δ*npx* (Δ*npx,* white squares), or Δ*npx:c* (Δ*npx:c,* blue triangles) were grown overnight in A) human amniotic fluid or B) mouse amniotic fluid and cell viability (Log_10_ CFU/mL) was determined by quantitative culture techniques. The Δ*npx* mutant was attenuated in cell viability in human and mouse amniotic fluid, a result that was reversed by genetic complementation *in trans*. ***P<0.001, ****P<0.0001, one-way ANOVA with Tukey’s multiple comparisons *post hoc* test.

**Supplemental Figure 2.**
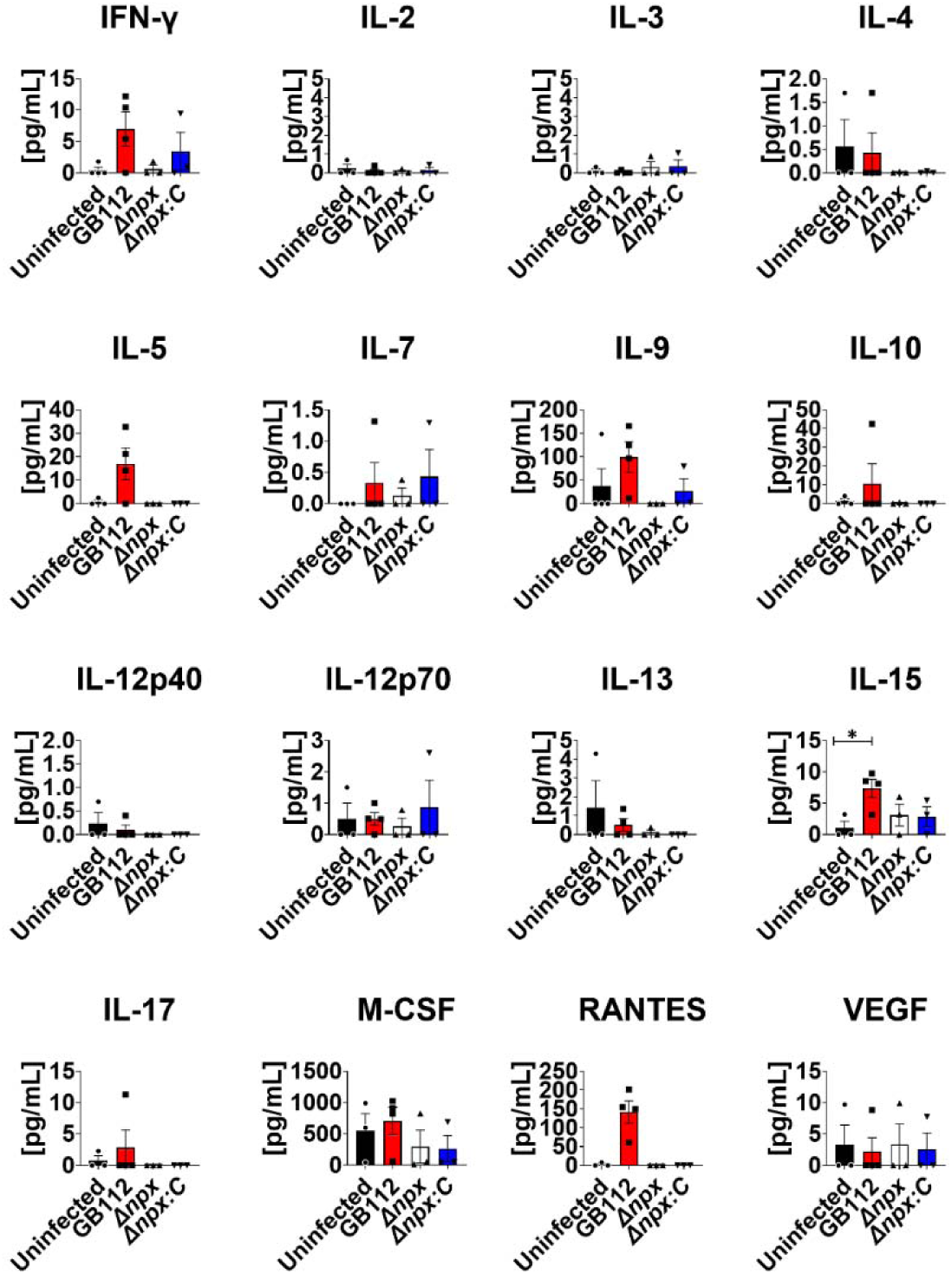
Analysis of cytokine production in uterus tissue in response to GBS infection. Multiplex cytokine analyses of uterine tissues after ascending vaginal infection with wild-type GB112 (GB112, red bars), Δ*npx* isogenic mutant (Δ*npx*, white bars), or the isogenic Δ*npx* complemented derivative (Δ*npx:c*, blue bars), as well as uninfected controls (UI, black bars). Uterine tissues were collected from pregnant mice on embryonic day E15.5, two days post- vaginal infection with GBS. Graphs indicate quantification of IFN-γ, IL-2, IL-3, IL-4, IL-5, IL-7, IL-9, IL-10, IL-12p40, IL-12p70, IL-13, IL-15, IL-17, M-CSF, RANTES, and VEGF levels. Bars indicate mean values +/- standard error mean with individual data points representing results from uterus tissues from individual dams. *P<0.05, one-way ANOVA with Tukey’s *post hoc* multiple comparisons test.

**Supplemental Figure 3.**
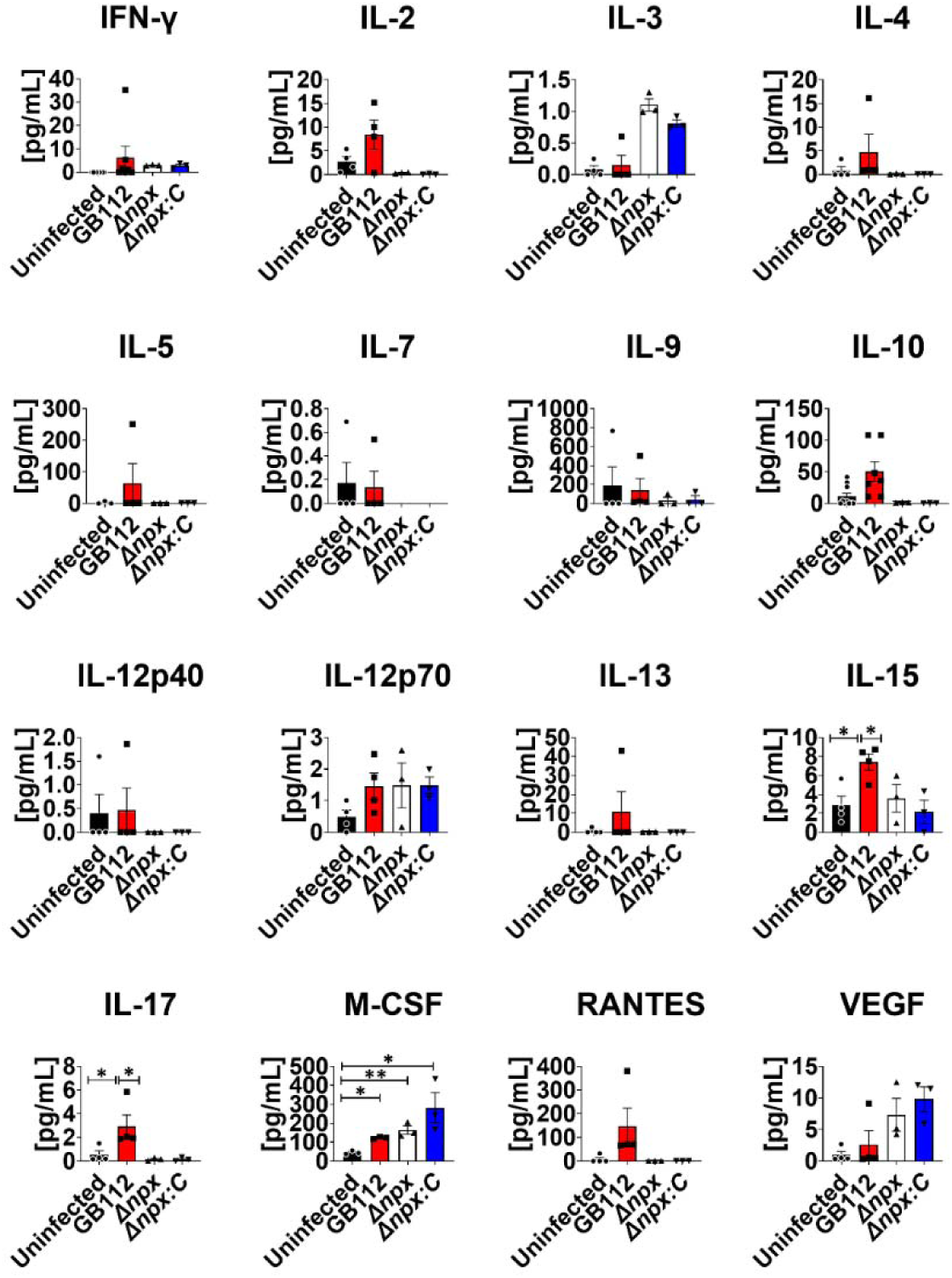
Analysis of cytokine production in decidua tissue in response to GBS infection. Multiplex cytokine analyses of decidual tissues after ascending vaginal infection with wild-type GB112 (GB112, red bars), Δ*npx* isogenic mutant (Δ*npx*, white bars), or the isogenic Δ*npx* complemented derivative (Δ*npx:c*, blue bars), as well as uninfected controls (UI, black bars). Decidual tissues were collected from pregnant mice on embryonic day E15.5, two days post- vaginal infection with GBS. Graphs indicate quantification of IFN-γ, IL-2, IL-3, IL-4, IL-5, IL-7, IL-9, IL-10, IL-12p40, IL-12p70, IL-13, IL-15, IL-17, M-CSF, RANTES, and VEGF levels. Bars indicate mean values +/- standard error mean with individual data points representing results from decidual tissues from individual dams. *P<0.05, **P<0.01, one-way ANOVA with Tukey’s *post hoc* multiple comparisons test.

**Supplemental Figure 4.**
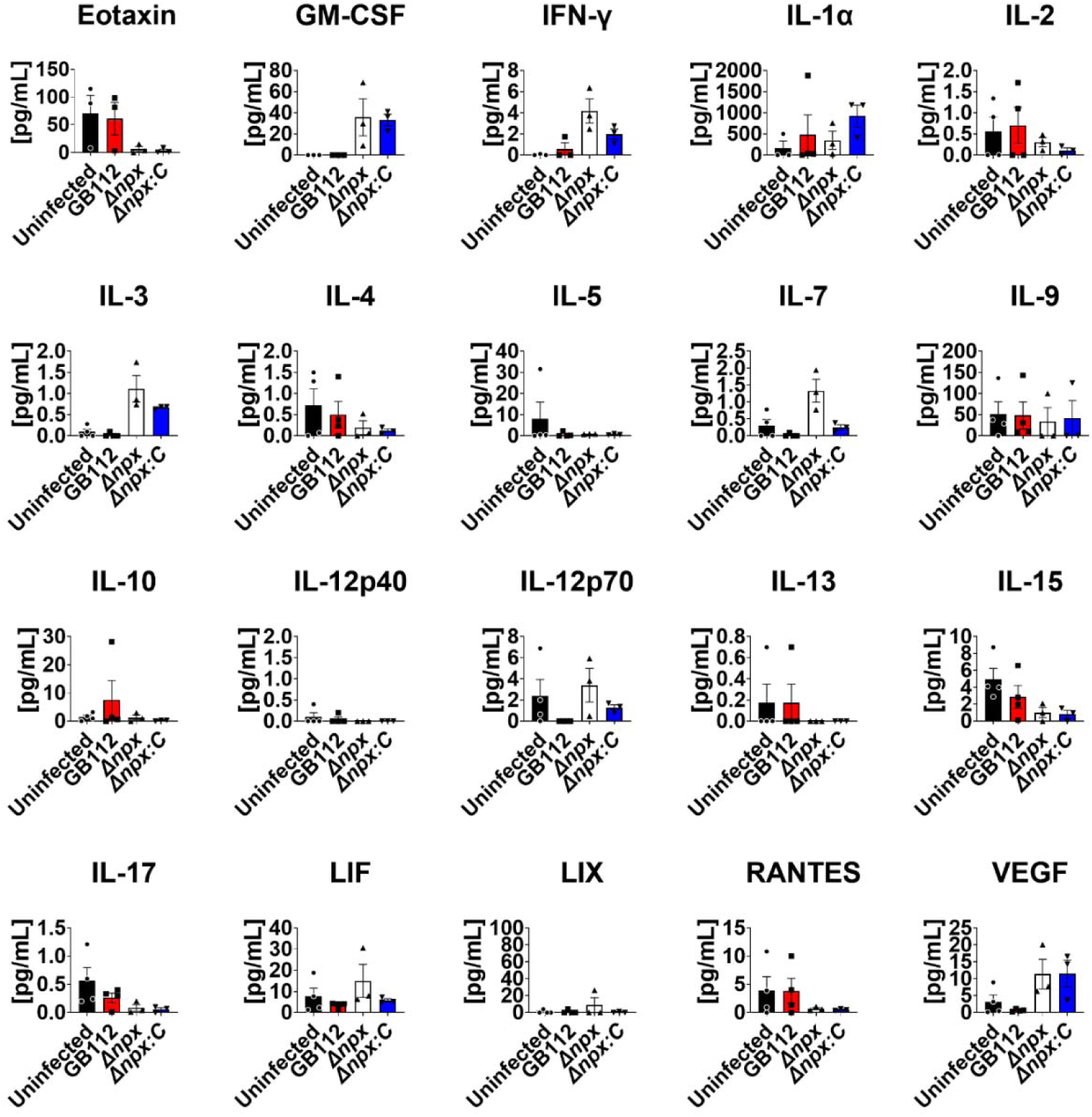
Analysis of cytokine production in placenta tissue in response to GBS infection. Multiplex cytokine analyses of placental tissues after ascending vaginal infection with wild-type GB112 (GB112, red bars), Δ*npx* isogenic mutant (Δ*npx*, white bars), or the isogenic Δ*npx* complemented derivative (Δ*npx:c*, blue bars), as well as uninfected controls (UI, black bars). Placental tissues were collected from pregnant mice on embryonic day E15.5, two days post- vaginal infection with GBS. Graphs indicate quantification of Eotaxin, GM-CSF, IFN-γ, IL- 1α, IL-2, IL-3, IL-4, IL-5, IL-7, IL-9, IL-10, IL-12p40, IL-12p70, IL-13, IL-15, IL-17, LIF, LIX, RANTES, and VEGF levels. Bars indicate mean values +/- standard error mean with individual data points representing results from decidual tissues from individual dams.

**Supplemental Figure 5.**
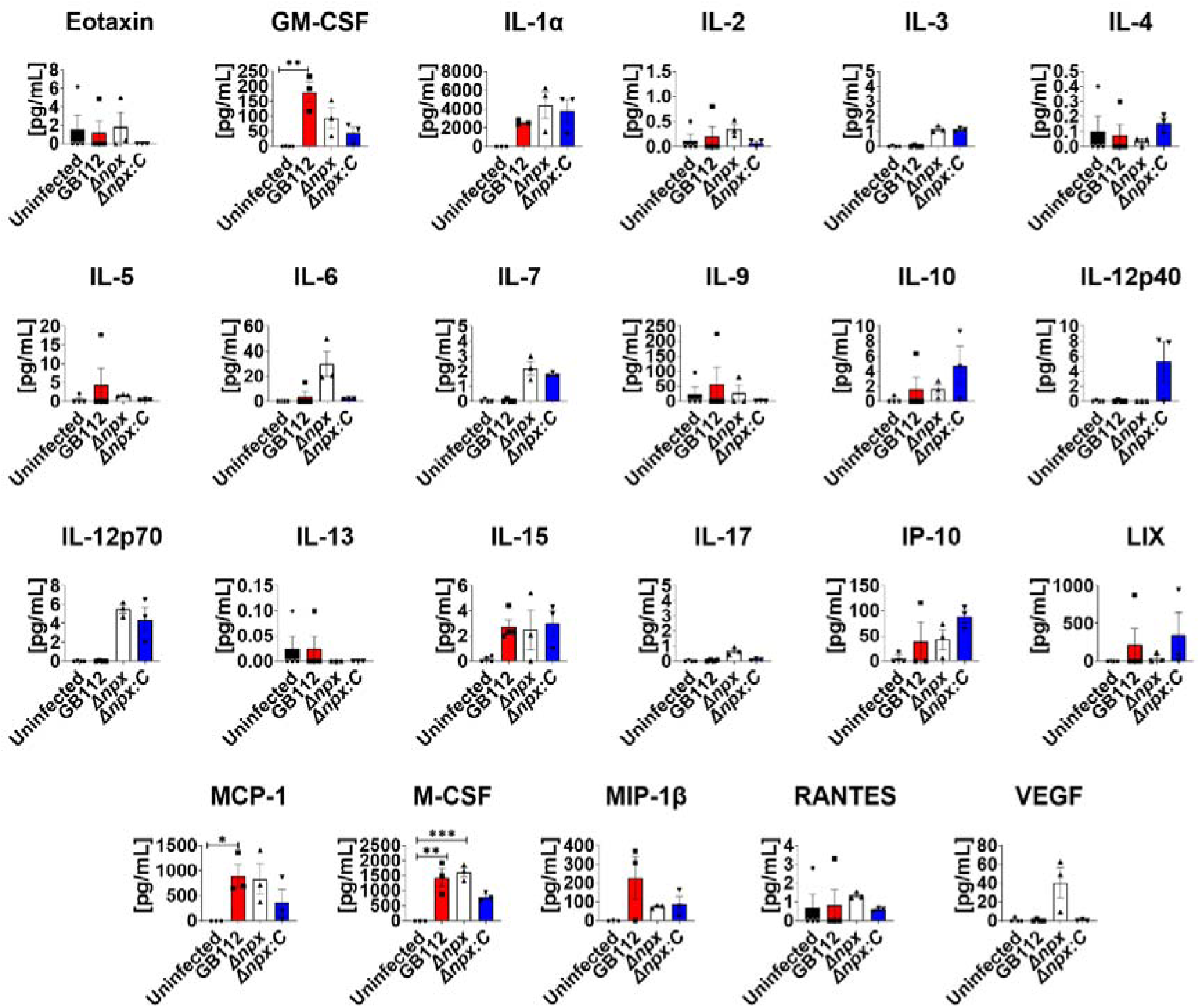
Analysis of cytokine production in amnion tissue in response to GBS infection. Multiplex cytokine analyses of placental tissues after ascending vaginal infection with wild-type GB112 (GB112, red bars), Δ*npx* isogenic mutant (Δ*npx*, white bars), or the isogenic Δ*npx* complemented derivative (Δ*npx:c*, blue bars), as well as uninfected controls (UI, black bars). Amnion tissues were collected from pregnant mice on embryonic day E15.5, two days post- vaginal infection with GBS. Graphs indicate quantification of Eotaxin, GM-CSF, IL-1α, IL-2, IL-3, IL-4, IL-5, IL-6, IL-7, IL-9, IL-10, IL-12p40, IL-12p70, IL-13, IL-15, IL-17, IP-10, LIX, MCP-1, M-CSF, MIP-1β, RANTES, and VEGF levels. Bars indicate mean values +/- standard error mean with individual data points representing results from amnion tissues from individual dams. *P<0.05, **P<0.01, ***P<0.001, one-way ANOVA with Tukey’s *post hoc* multiple comparisons test.

**Supplemental Figure 6.**
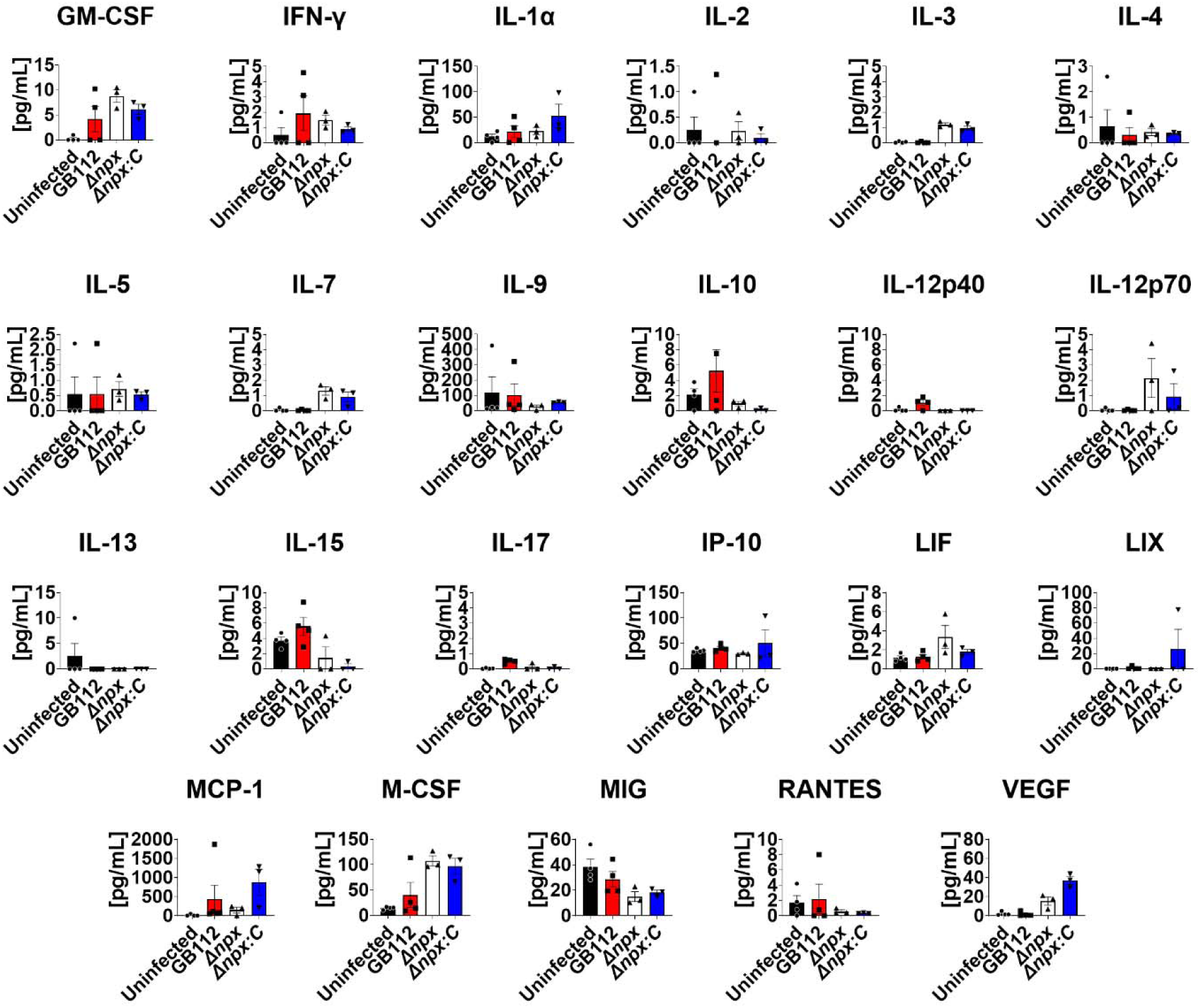
Analysis of cytokine production in fetus tissue in response to GBS infection. Multiplex cytokine analyses of fetal tissues after ascending vaginal infection with wild-type GB112 (GB112, red bars), Δ*npx* isogenic mutant (Δ*npx*, white bars), or the isogenic Δ*npx* complemented derivative (Δ*npx:c*, blue bars), as well as uninfected controls (UI, black bars). Fetal tissues were collected from pregnant mice on embryonic day E15.5, two days post-vaginal infection with GBS. Graphs indicate quantification of GM-CSF, IFN-γ, IL-1α, IL-2, IL-3, IL-4, IL-5, IL-7, IL-9, IL-10, IL-12p40, IL-12p70, IL-13, IL-15, IL-17, IP-10, LIF, LIX, MCP-1, M-CSF, MIG, RANTES, and VEGF levels. Bars indicate mean values +/- standard error mean with individual data points representing results from fetal tissues from individual dams.

## Notes

### Competing Interest Statement

The authors have declared no competing interest.

